# Threshold-Free Neural Network Models for Swim Bout Detection of Larval Zebrafish

**DOI:** 10.64898/2026.01.11.698899

**Authors:** Haochen Wang, Grant Kroeschell, Yunlong Zhu, Jeff S. Mumm, Ji Yi

## Abstract

Accurate detection of swim bouts is essential for decoding motor behavior in larval zebrafish, yet conventional threshold-based methods rely on subjective cutoffs and per-experiment tuning. To address this limitation, we developed threshold-free deep learning models operating on tail-curvature time series: an offline model for *post hoc* analysis and an online model for real-time detection. Both improved precision over threshold-based methods, particularly for low-amplitude bouts. Using the offline model, we show that a head-fixed, closed-loop preparation largely preserves naturalistic swimming kinematics. Repeated trials reduced bout frequency and increased interbout interval, whereas genetic background primarily affects bout duration and maximum amplitude; notably, casper larvae lacked the late-trial decline observed in wildtype and nacre. We further show that detector choice can alter inferred behavioral outcomes. Together, these results establish a high-throughput framework for robust and reproducible analysis of zebrafish swimming behavior across experimental conditions.

## Introduction

The brain evaluates sensory stimuli to generate and refine behavioral responses, and decoding this sensorimotor transformation is a central goal of neuroscience. The larval zebrafish (*Danio rerio*) is an established model for such studies, offering a vertebrate central nervous system amenable to high-throughput experimentation and system-wide neural imaging, with findings that can inform translational biomedical research^1–8^. To investigate the neural circuitry governing sensorimotor behavior, a common paradigm immobilizes the larval head for neural activity recording under a microscope while leaving the tail free to move, so that behavior can be tracked in response to controlled sensory stimuli^9^.

When analyzing the motor output of larval zebrafish, a fundamental step is the accurate detection of swim bouts - discrete, oscillatory tail movements separated by periods of quiescence^9, 10^. Conventionally, bouts are detected either by manual annotation or, more commonly, by applying a threshold to kinematic parameters such as tail angle, head orientation, or body position^11–22^. Although widely adopted within established analysis packages^15, 23, 24^, the threshold-based approach carries an intrinsic limitation: reliance on an arbitrary, user-defined cutoff. A low threshold risks misclassifying noise as a genuine bout, whereas a high threshold systematically misses subtle, low-amplitude events. Even when optimized, the threshold still requires per-experiment tuning that can undermine both reproducibility and the biological interpretation of the data. Deep learning and probabilistic methods have recently been adopted at other stages of the zebrafish behavioral analysis pipeline: markerless networks estimate pose directly from video^25–27^, and probabilistic models resolve the structure of already-segmented bouts^24, 28, 29^. The intervening step – segmenting bouts from the continuous tracking data – still relies on a user-defined threshold even within these pipelines.

Here, we address this gap with a threshold-free deep learning framework for bout detection. The core model couples a convolutional neural network (CNN) with a bidirectional long short-term memory (BiLSTM) layer, operates directly on the tail-curvature time series (the temporal derivative of Stytra’s tail_sum; **Methods**), and was trained on video-annotated ground truth. Positioned upstream of existing deep learning tracking and probabilistic bout-classification pipelines, it removes the user-defined cutoff while remaining accurate across signal regimes, with its largest gains over threshold-based detection on low-amplitude bouts that approach background noise. We validate the framework with anesthetized and hyperactive (*adgrl3.1* G0 knockout) controls, where the model recovers the expected absence of locomotion and the known hyperactive phenotype, and we show that our head-fixed closed-loop platform largely preserves naturalistic bout kinematics. Applying the validated framework to wildtype AB (WT), nacre (NCR), and casper (CSP) larvae, we find that prior trial experience and genetic background exert dissociable effects on swimming – trial experience broadly modulating bout kinematics while strain effects concentrate in bout duration and maximum amplitude – and that CSP larvae follows a late-trial trajectory distinct from WT and NCR cohorts. Finally, comparing the model with a benchmark threshold detector, we find that the two agree on the principal effects yet reach significance on different finer-grained comparisons, indicating that the choice of detector can itself shape the biological conclusions drawn. Together, these results provide a reproducible, high-throughput, threshold-free approach to bout detection that enables rigorous behavioral analysis across the genotypes and recording paradigms examined here.

## Results

### Behavioral Assays that Robustly Elicit Bouts for Automated Data Acquisition

The optomotor response (OMR) is a reflexive behavior in which animals – including insects^30^, fish^31^, and humans^32^ – move to stabilize their position relative to perceived optic flow. In zebrafish larvae, moving grating OMR assays reliably elicit swim bouts^31, 33^, so we adopted this paradigm to generate a large bout dataset efficiently.

We performed OMR assays in two configurations (**Figure 1A-B, Figure S1**; **Methods**). For free-swimming 6-7 days post-fertilization (dpf) larvae, a high-speed camera recorded tail kinematics at 200 Hz, which we tracked with the open-source software Stytra^34^, previously used for closed-loop zebrafish behavior^20, 35–37^. Stytra reduces tail curvature to a single scalar, a compact representation also adopted by other zebrafish tracking systems^21^. Retaining this tail_sum (*θ*) notation, we applied a temporal derivative to suppress slow baseline drift, such as the low frequency swing introduced by optic-table vibration, and isolate genuine tail motion (**Figure S5A; Methods**). A custom PsychoPy^38^ script projected moving red and black gratings – a robust OMR stimulus for zebrafish^39, 40^ – onto a diffusion pad beneath the fish, synchronized with tracking (**Figure S2; Table S1**). For head-fixed larvae, we mounted fish in agarose with the eyes and tail left free to move^41–44^ (**Figure 1C**; **Figure S3-S4**). Visual stimulation ran in two modes: in open-loop (OL), the grating moved at a constant, preset speed, decoupling sensory input from motor output; in closed-loop (CL), grating speed updated in real time from the fish’s intended motion, providing visual feedback (**Figure 1D**).

**Figure 1.**
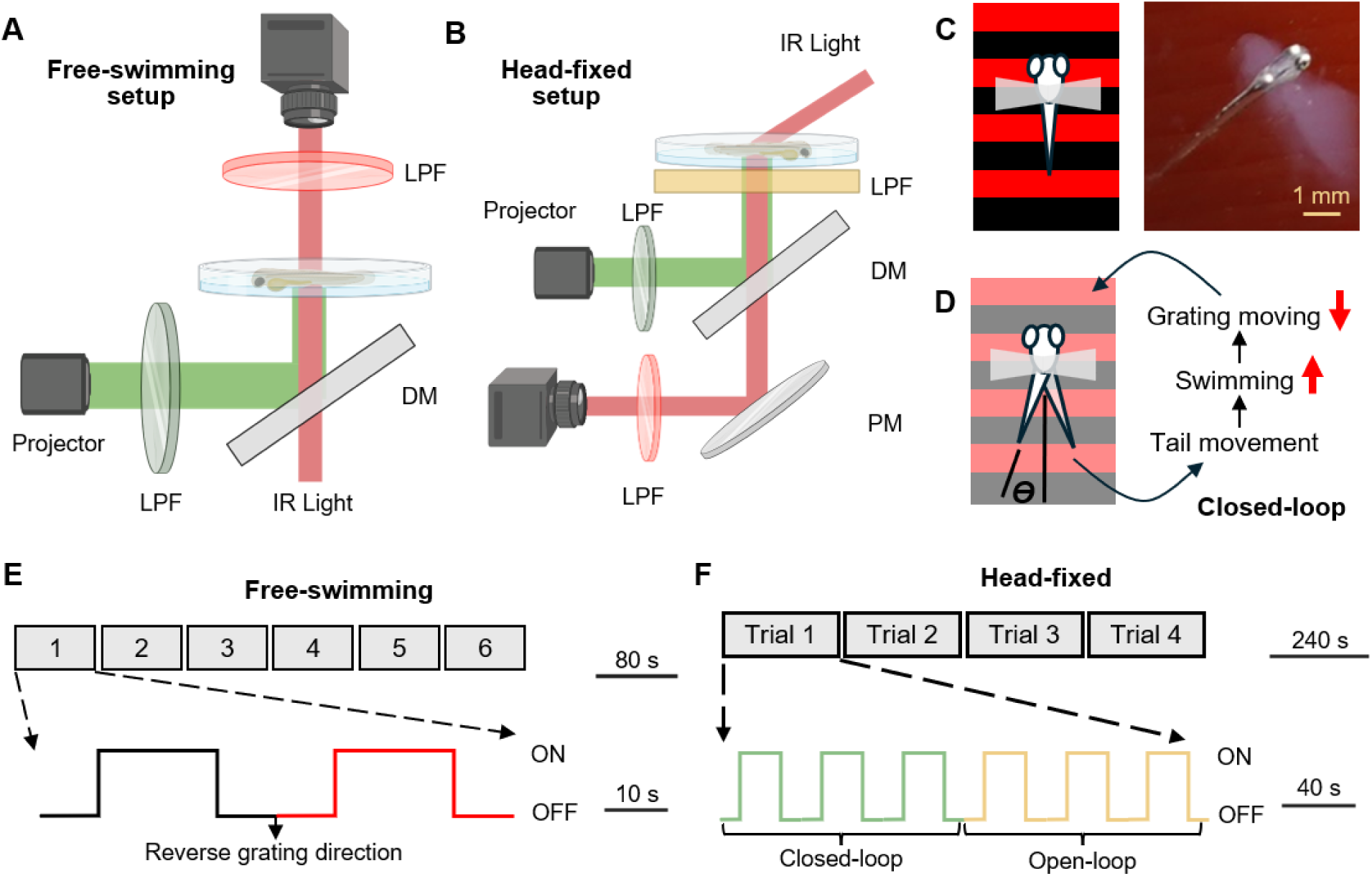
Experimental setups and behavioral paradigms for data collection. **(A) Free-swimming setup.** A top-down high-speed camera (200 Hz) images a single larva under infrared (IR) illumination from below, while the visual stimulus is projected through a long-pass filter (green LPF) to create black-on-red grating pattern onto a silicone diffusion pad supporting the 35 mm Petri dish. A cold (dichroic) mirror (DM) separates the IR imaging path from the visible stimulus path, and a long-pass filter (red LPF) blocks projector light so that only the IR-illuminated arena reaches the camera. **(B) Compact head-fixed closed-loop setup.** A folded optical path comprising a cold mirror (DM) and a plane mirror (PM) relays IR light for tail tracking, while the visible grating stimulus is reflected by the cold mirror onto a weighing-paper screen. Green and red LPFs are the same as in **Panel A**. The yellow LPF absorbs stray light from the microscope. **(C) Moving-grating stimulus schematic beneath a head-fixed larva**. The photo shows a larva in the agarose mount with its eyes and tail freed. During the movement (ON) phase, the grating drifts tail-to-head to evoke forward swimming via the optomotor response. **(D) Closed-loop feedback schematic**. Stytra converts the tracked tail signal (tail_sum, *θ*) into the larva’s intended forward velocity and reduces the grating speed accordingly, simulating a free-swimming experience. **(E) Free-swimming OMR protocol.** A 480 s session comprising six trials; each trial consists of two “20 s grating movement – 20 s pause” (ON/OFF) cycles, with the grating direction reversed between the two cycles. **(F) Head-fixed OMR protocol.** A 960 s session of four identical trials; each trial comprises three closed-loop (CL) cycles followed by three open-loop (OL) cycles of “20 s ON – 20 s OFF”. Panels A and B were created with BioRender.

Every stimulus cycle comprised 20 s of grating movement (ON) followed by 20 s of stasis (OFF). Free-swimming larvae completed six trials with alternating grating directions (**Figure 1E**). Head-fixed larvae completed four trials, each with three CL cycles followed by three OL cycles, the grating moving tail-to-head during ON (**Figure 1F**). Alternating CL and OL cycles diversified the evoked behaviors, enriching the dataset^31^.

### A Context-Aware Architecture for Threshold-Free Bout Detection

Because supervised training depends on reliable labels, we first annotated swim bouts in 14 randomly chosen sessions (N = 14 fish, one fish per session) spanning varied lighting conditions to challenge the model (**Figure S4**), with 8 sessions as the training set and 6 held-out sessions as the testing set. Each time point in the recorded tail traces was labeled as bout (1) or no-bout (0) through a two-stage procedure: an initial coarse pass over the synchronized video, followed by onset and offset refinement on the tail_sum (*θ*) signal (**Figure 2A**; **Figure S5B-C**). No amplitude or duration cutoff was imposed during annotation, ensuring that low-amplitude bouts were not excluded by construction. Inter- and intra-rater κ values and percent agreement are summarized in **Table S2**. Because bout labels represented only 14.2% of the training set and 16.1% of the testing set, model selection during 8-fold training and validation was guided by F1 score rather than accuracy, which would be less informative under this class imbalance (**Figure S6**).

**Figure 2.**
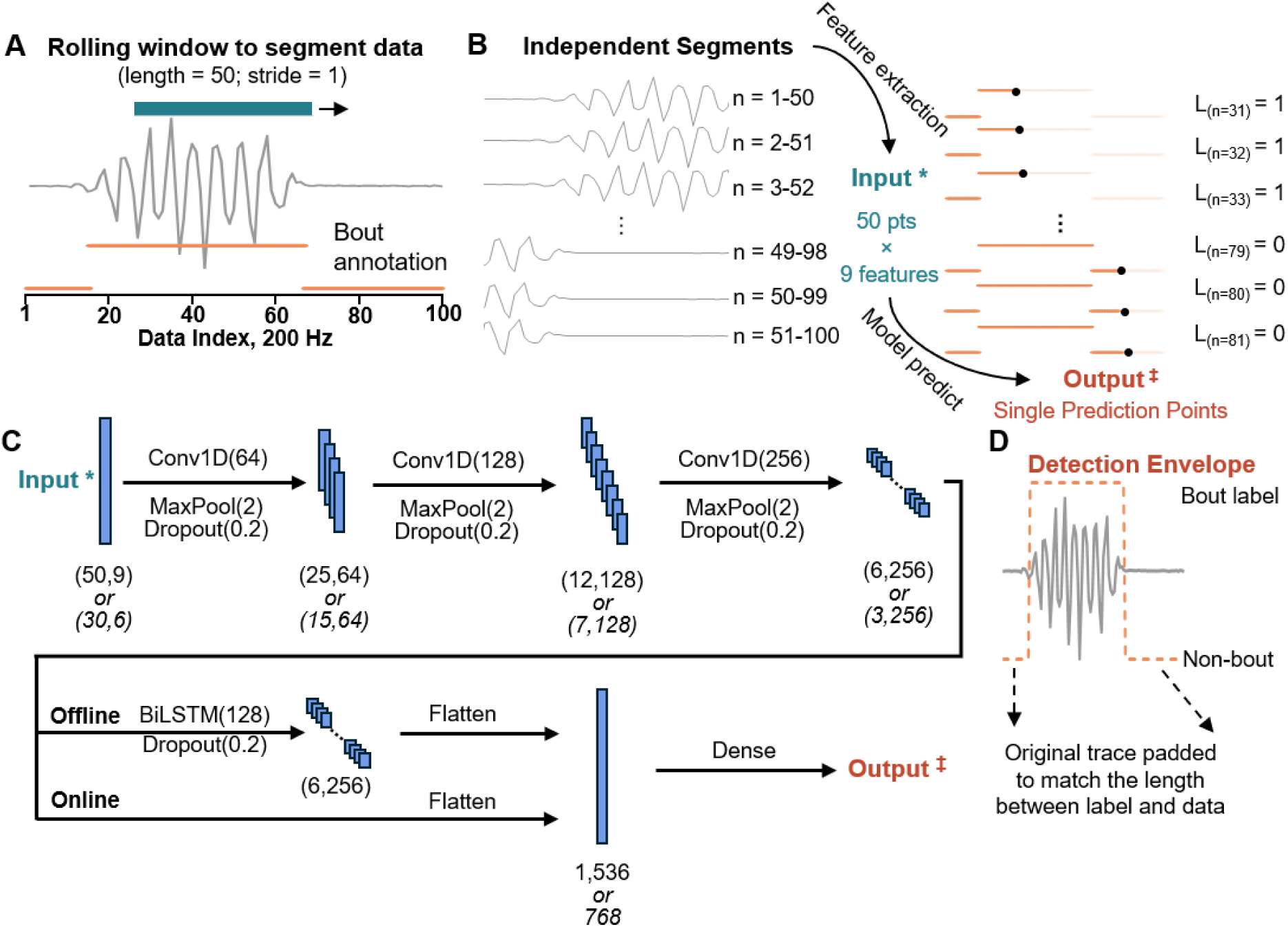
Data segmentation, inference, and architecture of the threshold-free bout-detection model. (A) Data segmentation. A 50-datapoint window (dark teal bar) slides one timepoint at a time along the entire tail_sum (*θ*) time series. Manually annotated training and testing sets supply a positive (bout) label at each timepoint where a bout occurs (orange bout envelope). **(B) Left and center panels:** From each window position a 50 × 9 feature matrix is extracted and passed to the model as input. **Right panel:** The model outputs a bout probability for the 31^st^ timepoint of the window; a 0.5 cutoff converts this into a binary label (1 = bout, 0 = no-bout). Successive calls concatenate into the continuous detection envelope for the whole dataset shown in **Panel D**. **(C) Offline and online neural-network architectures.** The offline model is used for *post hoc* data processing, and the online model for real-time bout detection. Both share three hierarchical one-dimensional convolutional blocks that extract features, each comprising a Conv1D layer (channels in parentheses), a Max Pooling layer (MaxPool; pooling size in parentheses), and a Dropout layer (rate in parentheses). The offline model adds a 128-unit bidirectional LSTM (BiLSTM) block that integrates the full window in both temporal directions to capture temporal dependencies within the segment, whereas the online model omits it for speed. Both terminate in a Flatten layer followed by Dense (fully connected) layers that produce the final score, with segments scoring above 0.5 classified as positive (bout) and the rest as negative (no-bout). Output shapes are labeled below each layer (top row = offline model; bottom row, where applicable = online model). **(D) Inference schematic**. The raw trace is zero-padded so that the detection-envelope length matches the original dataset. As the segmentation window advances, the per-segment labels concatenate into a sequential detection envelope.

Bout kinematics vary both within and across larvae^45, 46^, and low-amplitude tail movements can resemble background noise, so labels cannot be assigned reliably from instantaneous signal values alone. We therefore designed the model to be context-aware: a 50-point window slides along the tail_sum trace, and the model predicts the label of the window’s 31st point from the full surrounding context (**Figure 2A-B**). From each window we computed nine features drawn from three sources (**Table S3**): the raw Stytra tracking signal^34^, seven time-series statistics adapted from prior signal-analysis work^34, 47–50^, and a theory-driven feature based on known zebrafish kinematics (the 20-40 Hz band-pass filtered tail_sum power)^51^. A permutation feature importance analysis confirmed that each retained feature contributed to prediction^52^ (**Table S4-S5**).

The resulting feature matrix is passed through three convolutional blocks, followed by the BiLSTM layer and a final dense layer with a 0.5 cutoff that returns a bout or no-bout call for each timepoint (**Figure 2C**). Applying this classifier across the full trace produces a continuous “detection envelope” (**Figure 2D**). The model’s parameters were set to match the characteristic time scale of larval swimming: a bout lasts a few tenths of a second (0.3-0.45 s) and beats at roughly 20-25 Hz^53, 54^. At 200 Hz, the 50-point (250 ms) window is therefore comparable to a single bout and spans several tail-beat cycles. Within this window, the convolutional kernels capture local periodic structure, whereas the bidirectional long short-term memory (BiLSTM) layer integrates the full window in both temporal directions. This full-window integration allows the model to evaluate whether a low-amplitude oscillation persists like a bout or dissipates like noise, rather than relying on instantaneous amplitude. We refer to this *post hoc* analysis configuration as the offline model.

For real-time, closed-loop experiments, we also trained a streamlined online model with a shorter feature window and no BiLSTM to improve inference speed (**Figure 2C**; **Figure S7**), raising the combined real-time tracking and detection rate from ∼50 Hz with the offline model to ∼150 Hz.

### Models Show Robust Detection Across Bout Amplitudes

We first assessed overall model performance on the held-out testing set (N = 6 fish). The offline model reached an area under the receiver operating characteristic (ROC) curve of 0.9945 (**Figure 3A**), and its precision-recall curve gave an average precision of 0.9794 (**Figure 3B**). The F1 score was maximized at a classification cutoff of 0.4888, confirming that the standard 0.5 decision threshold we use was well placed. The online model performed comparably (ROC AUC 0.9900, average precision 0.9724, best F1 at a 0.4994 cutoff; **Figure S8**).

**Figure 3.**
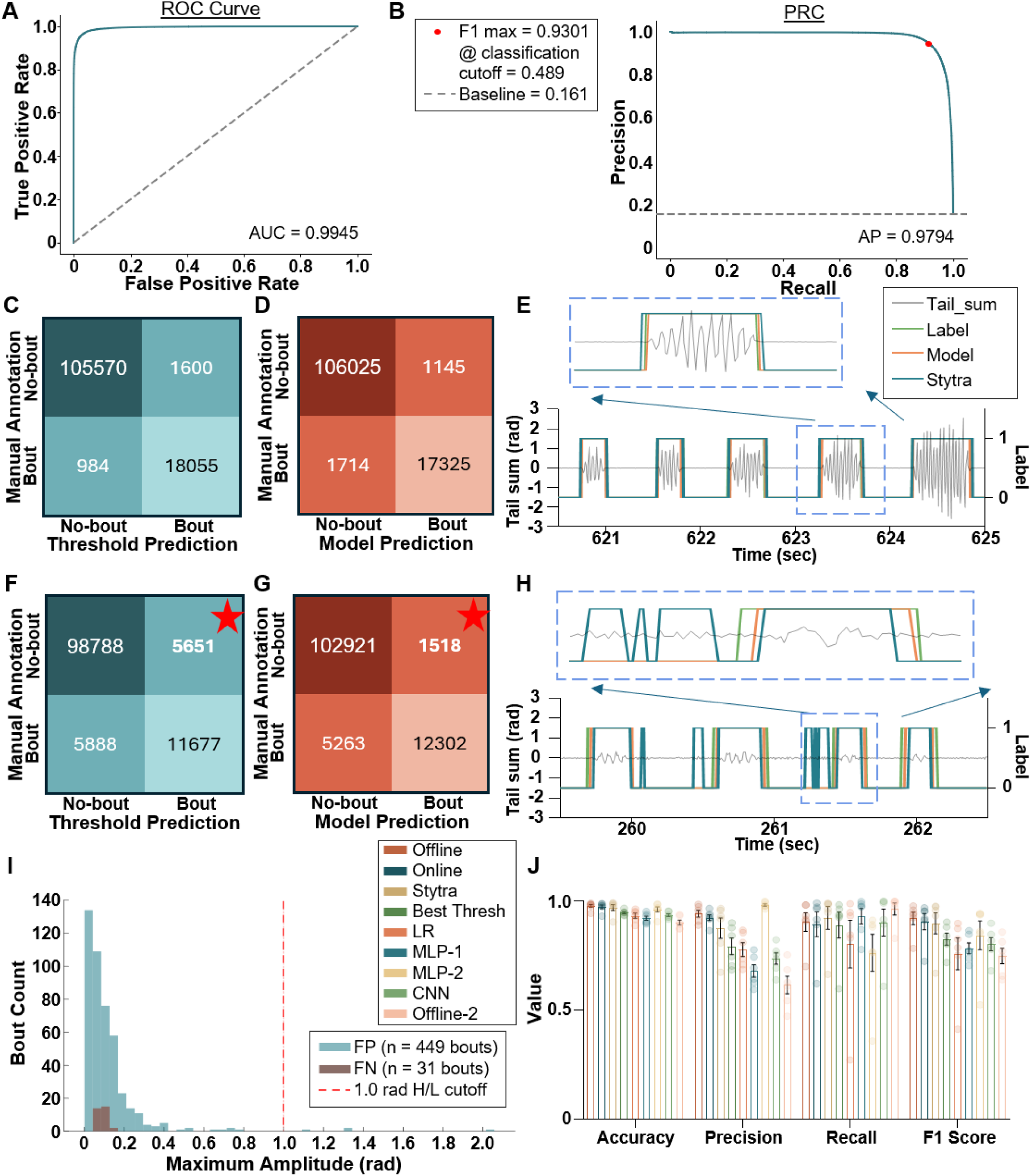
Evaluation of bout-detection performance. Head-fixed testing set (N = 6 fish, 6-7 dpf, AB wild-type). **(A) Receiver-operating-characteristic (ROC) curve of the offline model on the testing set.** The area under the curve (AUC) = 0.9945. **(B) Precision-recall curve of the offline model.** Average precision (AP) = 0.9794 (prevalence baseline = 0.161). The F1 score is maximized at a classification cutoff of 0.4888, close to the fixed 0.5 cutoff used throughout. **(C-E) A representative high-amplitude session (56.12% of bouts exceed a 1.0 rad peak-to-peak amplitude), where both detectors perform comparably.** Threshold: accuracy 0.9795, precision 0.9186, recall 0.9483, F1 0.9332; offline: accuracy 0.9773, precision 0.9380, recall 0.9100, F1 0.9238. **(E)** shows comparison at 620.5-625 s, subpanel zooms to 623-624 s, revealing only minor onset/offset shifts. **(F-H) A representative low-amplitude session (7.87% of bouts exceed 1.0 rad), where the offline model surpasses the threshold detector on every metric.** Threshold: accuracy 0.9054, precision 0.6739, recall 0.6648, F1 0.6693; offline: accuracy 0.9444, precision 0.8902, recall 0.7004, F1 0.7839. **(H)** shows comparison at 259.5-262.5 s, subpanel zooms to 261.2-261.7 s, where the threshold detector flags low-amplitude fluctuations as bouts. **(I) Peak-to-peak amplitude of offline-model false positive (FP) and false negative (FN) on the testing set**. The red dashed line marks 1.0 rad. A bout is scored as detected (a true positive) when the offline model emits at least one bout label within a manually labeled bout. 445 of 449 FP and all 31 FN have amplitude below 1.0 rad. **(J) Accuracy, precision, recall, and F1 (bars = mean ± SEM across the N = 6 testing fish) for the detectors compared**. Offline (three CNN blocks + BiLSTM), Online (three CNN blocks), Stytra (10-point rolling standard deviation, threshold 0.050), Best Threshold (10-point moving average of 20-40 Hz band-pass-filtered tail_sum power, 4th-order Butterworth, single global threshold; Pow10S), LR (logistic regression), MLP-1 (perceptron on per-window feature summaries that summarize the temporal component), MLP-2 (perceptron on single-timepoint features that remove the temporal component), CNN (single convolutional layer), and Offline-2 (offline architecture on the six raw tail-segment angles). Group-mean F1: Offline 0.9212, Online 0.9057, Stytra 0.8983, Best Threshold 0.8262, LR 0.7584, MLP-1 0.7832, MLP-2 0.8435, CNN 0.8028, Offline-2 0.7482. *Traces (**C, H**): grey, tail_sum tracking from Stytra; green envelopes, manually labeled bouts; orange envelopes, offline-model detections; dark teal envelopes, threshold-method detections. See the Methods section for definitions of tail_sum and the bout-detection operations*.

Next, we compared the offline model with the threshold-based method built into Stytra^34^, run at its default threshold of 0.050. Across the testing set, the offline model achieved an accuracy of 0.9782, precision of 0.9426, recall of 0.9044, and an F1 score of 0.9212 (**Table S6**). The online model degraded only marginally – accuracy 0.9742 (0.0040 lower) and F1 0.9057 (0.0155 lower) – indicating that, when inference time is not limiting, the offline model retaining the BiLSTM layer is preferred (**Table S6**). Examining two sessions where the offline had the lowest F1 score, we found it matched the threshold method on one and produced far fewer false positives on the other (**Figure 3C-H**). On the first session, both methods performed well, with comparable accuracy (0.9773 offline vs 0.9795 threshold; **Figure 3C-D**); the offline model traded higher precision (0.9380 vs 0.9186) for lower recall (0.9100 vs 0.9483), and both agreed closely with manual annotation (**Figure 3E**). On the second session, the offline model surpassed the threshold method on every metric (**Figure 3F-G**): precision rose most, from 0.6739 to 0.8902, alongside gains in accuracy (0.9054 to 0.9444), recall (0.6648 to 0.7004), and F1 (0.6693 to 0.7839). Visual inspection showed that the threshold method flagged fluctuating noise as bouts, generating thousands of false positives that the context-aware model correctly ignored (**Figure 3H**).

To understand these failures, we examined the peak-to-peak amplitude (the maximum minus the minimum tail_sum within a bout) of every labeled bout in the testing set. In the two example sessions, 56.12% and 7.87% of bouts exceeded a peak-to-peak amplitude of 1.0 rad, and the offline model’s false positives fell below this value (**Figure 3I**; **Figure S9; Movie S1**). Within each of the six testing sessions, this amplitude showed no consistent temporal trend (**Figure S10**; **Table S7**), so it is a property of the session rather than of time within it. We therefore classified a session as high amplitude when more than half its bouts exceeded 1.0 rad and low amplitude otherwise.

Varying the Stytra threshold on the second, low amplitude session (where 7.87% of bouts exceeded 1.0 rad; **Figure 3F-G**) exposed the core limitation: low values (≤0.060) produced more false positives, while high values (≥ 0.065) missed the subtle onset and offset of low amplitude bouts (**Figure S11A**). A threshold of 0.060 outperformed the default 0.050 and was the best of all tested values (**Figure S11B-C**), showing that threshold-based detection requires per-experiment tuning. Using contextual information, the offline and online models exceeded the accuracy and F1 score of every tested threshold on this session.

To test whether a better-designed threshold could close the gap, we expanded the benchmark to 32 additional variants spanning four statistic families (rolling standard deviation, rolling mean, rolling median, and the moving average of the 20-40 Hz band-pass filtered tail_sum power), four window lengths (10, 50, 70, and 100 points; 50-500 ms at 200 Hz, from a fraction of a single bout to longer than one), and both global and per-session adaptive calibration. All variants were tuned on the training set and evaluated on the challenging low amplitude sessions of the testing set, except that per-session adaptive calibration was directly performed on the low amplitude sessions (**Figure S12A**).

On these challenging sessions, the highest-scoring variant was the 10-point moving average of the 20-40 Hz band-pass filtered tail_sum power with a threshold tuned per fish (Pow10T, F1 0.8239), followed by the per-fish tuning 10-point moving standard deviation (Std10T, F1 0.8127) and per-fish tuning 10-point moving mean (Mean10T, F1 0.8083), and then the filtered power with a single global threshold (Pow10S, F1 0.7864). All variants performed below the offline model (F1 0.8608), which itself trailed its full-test-set performance (0.9212) due to the greater difficulty of low-amplitude sessions. The last three also fell below the online model (0.8183).

Two features of this benchmark are notable. First, matching the window length to the mean bout duration did not improve performance: the 10-point window beat the duration-matched window across all four statistic families, so the model’s advantage is not an artifact of window misspecification. Second, per-session calibration – scaling each session’s threshold by the standard deviation of its non-bout baseline noise – improved on global calibration, but it requires session-level, manual labeling for the scaling sweep, which is the very labeling cost the deep learning model removes. For downstream analysis of larger datasets, we therefore adopted the best global variant – the 10-point moving average of the 20-40 Hz band-pass filtered tail_sum power (Pow10S) – which we refer to as the “best threshold method.”

On the full test set we further compared both models against simpler machine learning baselines: logistic regression on the flattened feature matrix (no nonlinear boundary; F1 0.7584); a multilayer perceptron on aggregated features (MLP-1, each feature summarized by its mean, standard deviation, minimum, maximum, and current value; F1 0.7832); a multilayer perceptron on instantaneous features only (MLP-2, the value at the current timestep for all features, removing temporal context; F1 0.8435); a single convolutional layer (CNN; F1 0.8028); and the offline architecture run on the six raw tail segment angles (Offline-2; F1 0.7482). The offline (F1 0.9212) and online (F1 0.9057) models outperformed all of these (**Figure 3J**; **Figure S12B-E**).

Finally, to test generalizability beyond the head-fixed dataset, we evaluated both models on three labeled free-swimming sessions (N = 3 fish) recorded under varying lighting conditions and noise levels (**Figure S4**). Performance decreased relative to the head-fixed test set, but both the offline (F1 0.8911) and online (F1 0.8628) models consistently outperformed the native Stytra threshold method (F1 0.7559) and the best threshold method (F1 0.8900) across all sessions (**Table S8**). These results indicate that the models transfer across recording conditions without the need for per-experiment threshold tuning.

### The Offline Model Detects Known Strain Differences and Resolves Feedback-Dependent Bout Kinematics

We next asked whether the offline model could recover strain-dependent bout kinematics consistent with previous publications. Because *adgrl3.1^-/-^* larvae have been reported to swim further and spend more time immobile than wildtype^55, 56^, we generated *adgrl3.1* knockouts in eleven AB fish (N = 11 fish) and assayed them head-fixed (**Figure 1B-D**, **1F**) at 7 dpf as a positive control, using 55 wildtype AB fish at 6-7 dpf (N = 55 fish) as the baseline. By 2-way ANOVA on per-fish per-trial means (strain × trial) and *post hoc* Tukey’s test (α = 0.05), bout duration showed a significant strain effect during grating movement in both closed-loop (CL-ON, p = 0.0001) and open-loop (OL-ON, p = 0.0016) conditions, but not during grating pause (CL-OFF, p = 0.8603; OL-OFF, p = 0.3989). Pairwise comparisons showed that *adgrl3.1* KO fish produced longer bouts than wildtype in CL-ON Trial 2 (p = 0.0024) and Trial 4 (p = 0.0211) (**Figure 4A-D**). Interbout interval also showed a significant strain effect in all four conditions (CL-ON, p < 0.0001; CL-OFF, p = 0.0259; OL-ON, p = 0.0001; OL-OFF, p = 0.0310), with *adgrl3.1* KO fish resting longer between bouts (**Figure 4E-H**). These observations are consistent with previously reported behavioral differences in *adgrl3.1* KO larvae and demonstrate that the model can resolve strain-dependent differences in bout kinematics. Extending the comparison to additional parameters, we found strain effects in some or all conditions for bout frequency (CL-ON p = 0.0034, OL-ON p = 0.0221; n.s.: CL-OFF p = 0.4769 and OL-OFF p = 0.2666), tail beat frequency (CL-ON p = 0.0132, CL-OFF p = 0.0373, OL-OFF p = 0.0487; n.s.: OL-ON p = 0.0731), and maximum amplitude (all four conditions significant: CL-ON p = 0.0082, CL-OFF p = 0.0005, OL-ON p = 0.0026, OL-OFF p = 0.0030), with *adgrl3.1* KO fish generally lower than wildtype (**Figure S13**). These differences suggest additional features of the knockout phenotype that warrant future study.

**Figure 4.**
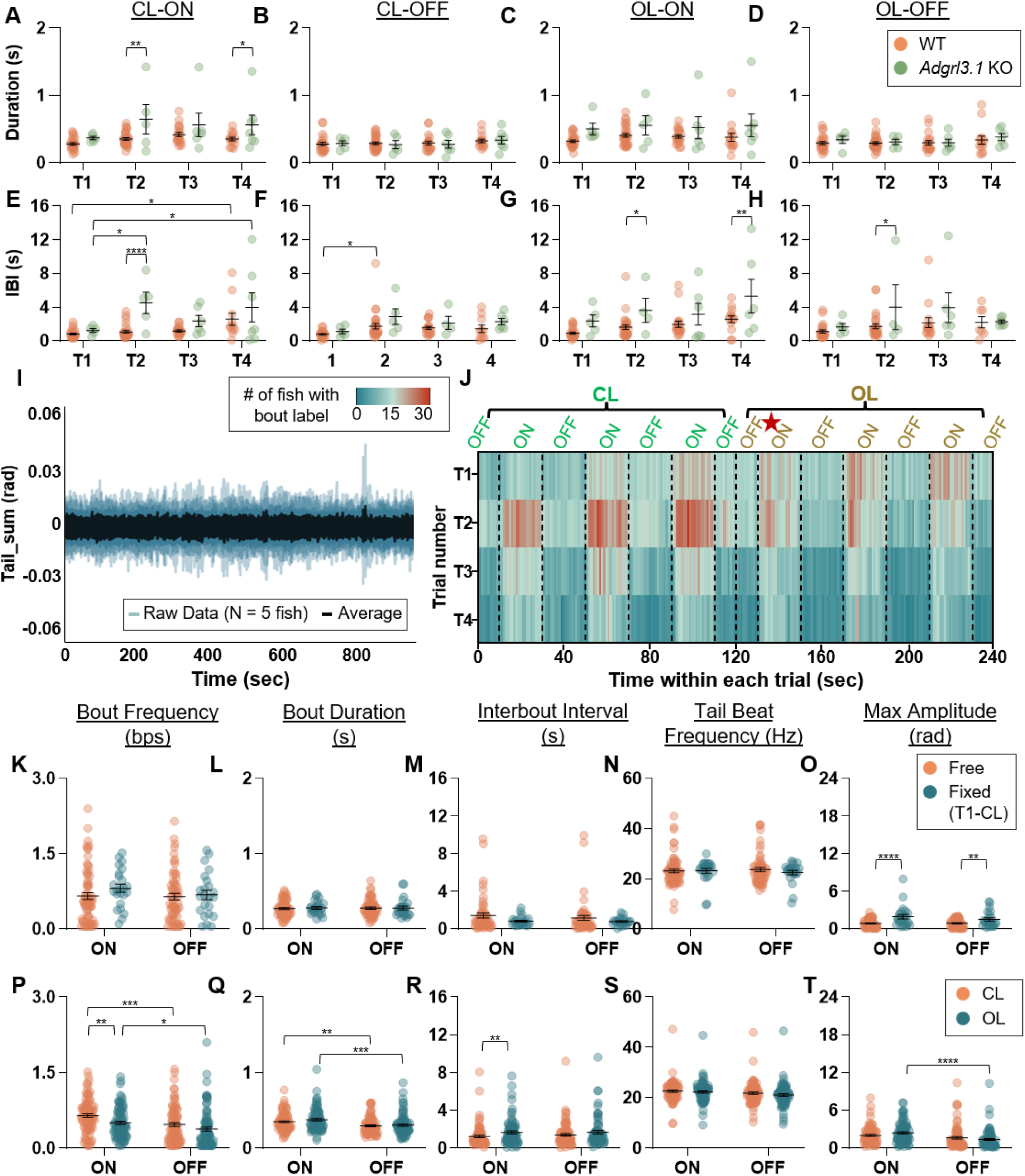
Modulation of bout kinematics by genotype and visual feedback in head-fixed and free-swimming larvae. Datasets by panel block: **A-H**, adgrl3.1 knockout (KO; N = 11, 7 dpf, AB, head-fixed) versus WT AB (N = 55, 6-7 dpf, head-fixed); **I**, tricaine-anesthetized larvae (N = 5; four at 12 dpf, one at 7 dpf, head-fixed); **J**, head-fixed WT AB (N = 55); **K-O**, free-swimming WT AB (N = 11, 6-7 dpf) versus head-fixed WT AB (N = 55, closed-loop Trial 1 only); **P-T**, head-fixed WT AB (N = 55), all four trials. Closed-loop feedback during recording was driven by the Stytra threshold detector; the bout labels analyzed here were generated *post hoc* by the offline model. Two-way ANOVA throughout; **A-H** use Tukey’s post-hoc, **K-T** use Fisher’s LSD. 95% CI represents the confidence interval for the difference of the mean in pairwise comparison; negative mean difference or CI means the first cohort is less than the second, vice versa. Bars = mean ± SEM; points = single fish. **(A-D) Bout duration of *adgrl3.1* KO versus WT at CL-ON, CL-OFF, OL-ON, OL-OFF**. A strain effect is present during grating movement (CL-ON p = 0.0001; OL-ON p = 0.0016) but not during the pause (CL-OFF p = 0.8603; OL-OFF p = 0.3989). **(A)** KO bouts are longer than WT at CL-ON Trial 2 (WT < KO: mean difference -0.2916, 95% CI -0.4769 to -0.1062, p = 0.0024) and Trial 4 (WT < KO: mean difference -0.2088, 95% CI -0.3855 to -0.03207, p = 0.0211). **(E-H) Interbout interval of *adgrl3.1* KO versus WT**. A strain effect is present in all four conditions (CL-ON p < 0.0001; CL-OFF p = 0.0259; OL-ON p = 0.0001; OL-OFF p = 0.0310), with KO larvae resting longer between bouts. All significant pairwise comparisons: **(E)** at CL-ON, KO longer than WT at Trial 2 (WT < KO: mean difference - 3.426, 95% CI -4.986 to -1.865, p < 0.0001), with interbout interval also lengthening across trials within each strain (WT T1 < T4: mean difference -1.749, 95% CI -3.291 to -0.2078, p = 0.0195; KO T1 < T2: mean difference -3.259, 95% CI -5.937 to -0.5819, p = 0.0104; KO T1 < T4: mean difference -2.731, 95% CI -5.210 to -0.2520, p = 0.0248); **(F)** at CL-OFF, WT T1 < T2 (mean difference -0.9651, 95% CI -1.893 to -0.03764, p = 0.0381); **(G)** at OL-ON, KO longer than WT at Trial 2 (WT < KO: mean difference -2.001, 95% CI -3.980 to -0.02175, p = 0.0476) and Trial 4 (WT < KO: mean difference -2.743, 95% CI -4.570 to -0.9155, p = 0.0037); **(H)** and at OL-OFF, KO longer than WT at Trial 2 (WT < KO: mean difference -2.266, 95% CI -4.454 to -0.07857, p = 0.0425). **(I) Aggregated tail_sum traces from five tricaine-anesthetized larvae (negative control)**. The offline model detects zero bouts in all five sessions. **(J) Heatmap of bout occurrences over time for the N = 55 head-fixed WT AB cohort (1 s bins; four trials as row blocks).** The red star marks the sharp activity drop at the onset of each trial’s first open-loop cycle. **(K-O) Free-swimming versus head-fixed WT AB (closed-loop Trial 1)** for bout frequency **(K)**, duration **(L)**, interbout interval **(M)**, tail-beat frequency **(N)**, and maximum amplitude **(O)**. Frequency, duration, interbout interval, and tail-beat frequency do not differ between groups (**K**: ON p = 0.1943, OFF p = 0.7840; **L**: ON p = 0.7385, OFF p = 0.9579; **M**: ON p = 0.1235, OFF p = 0.3400; **N**: ON p = 0.9666, OFF p = 0.3817). **(O)** Maximum amplitude is higher in head-fixed than free-swimming larvae during both grating movement (Fixed > Free: mean difference 1.088, 95% CI 0.6439-1.531, p < 0.0001) and pause (Fixed > Free: mean difference 0.6156, 95% CI 0.1603-1.071, p = 0.0083). **(P-T) Closed-loop versus open-loop within the head-fixed** cohort for frequency **(P)**, duration **(Q)**, interbout interval **(R)**, tail-beat frequency **(S)**, and maximum amplitude **(T)**. **(P)** Closed-loop raises bout frequency relative to open-loop during grating movement (CL-ON > OL-ON: mean difference 0.1473, 95% CI 0.04369-0.2508, p = 0.0055) but not during pause (p = 0.0964); grating movement raises frequency relative to pause within both feedback conditions (CL ON > OFF: mean difference 0.1792, 95% CI 0.07532-0.2831, p = 0.0008; OL ON > OFF: mean difference 0.1202, 95% CI 0.01636-0.2241, p = 0.0234). **(Q)** Duration is modulated by grating movement (CL ON > OFF: mean difference 0.05376, 95% CI 0.01538-0.09214, p = 0.0062; OL ON > OFF: mean difference 0.07286, 95% CI 0.03447- 0.1112, p = 0.0002) but not by feedback condition (CL vs OL: ON p = 0.1719, OFF p = 0.7004). **(R)** Interbout interval is shorter under closed-loop during grating movement (CL-ON < OL-ON: mean difference -0.4336, 95% CI -0.8453 to -0.02184, p = 0.0391) but not during pause (p = 0.1942). **(S)** Tail-beat frequency shows a main effect of grating phase (ON vs OFF p = 0.0342) with no closed- versus open-loop effect (CL vs OL p = 0.2302) and no significant pairwise difference (CL-ON vs CL-OFF p = 0.2020; OL-ON vs OL-OFF p = 0.0848). **(T)** Maximum amplitude is modulated by grating movement, most strongly under open-loop (OL ON > OFF: mean difference 1.067, 95% CI 0.5973-1.537, p < 0.0001; CL ON vs OFF p = 0.0952).

As a negative control, we recorded five anesthetized fish (tricaine-treated; N = 5 fish, one at 7 dpf and four at 12 dpf). The offline model detected no bouts in any session (**Figure 4I**).

We then examined the wildtype AB cohort (N = 55 fish). Bout activity increased from Trial 1 to Trial 2, and then declined steadily across later trials (**Figure 4J**). At the onset of the first open-loop cycle of each trial, swimming dropped sharply (**Figure 4J**, red star), consistent with larvae detecting the sensory mismatch when concordant visual feedback was removed^33, 57, 58^.

To test whether head mounting altered behavior, we compared head-fixed larvae during the closed-loop period of Trial 1 – before any exposure to open-loop cycles – with a free-swimming cohort (N = 11 WT AB fish, 6-7 dpf) by 2-way ANOVA and *post hoc* Fisher’s LSD test (α = 0.05, p-values are reported from Fisher). Bout frequency (ON p = 0.1943, OFF p = 0.7840), duration (ON p = 0.7385, OFF p = 0.9579), interbout interval (ON p = 0.1235, OFF p = 0.3400), and tail beat frequency (ON p = 0.9666, OFF p = 0.3817) did not differ between groups (**Figure 4K-N**). Maximum bout amplitude, however, was higher in head-fixed than in free-swimming fish during both grating movement (p < 0.0001) and pause (p = 0.0083) (**Figure 4O**).

Comparing closed-loop and open-loop conditions within the head-fixed cohort (N = 55, Trial 1 to Trial 4), by 2-way ANOVA and *post hoc* Fisher’s LSD test (α = 0.05, p-values are reported from Fisher unless stated otherwise), further showed how real-time feedback modulates the optomotor response. Closed-loop feedback increased bout frequency relative to open-loop during grating movement (CL-ON vs. OL-ON, p = 0.0055) but not during pause (CL-OFF vs. OL-OFF, p = 0.0964). Grating movement also increased bout frequency relative to pause within both closed-loop (p = 0.0008) and open-loop (p = 0.0234) conditions (**Figure 4P**). Bout duration was modulated by grating movement (ON vs. OFF within CL, p = 0.0062; within OL, p = 0.0002) but not by feedback condition (CL vs. OL: ON p = 0.1719, OFF p = 0.7004) (**Figure 4Q**). Interbout interval was shorter under closed-loop during grating movement (CL-ON vs. OL-ON, p = 0.0391) but not during pause (p = 0.1942) (**Figure 4R**). Tail beat frequency showed a grating ON versus OFF phase main effect (p = 0.0342; 2-way ANOVA), although no individual pairwise comparison was significant (CL p = 0.2020, OL p = 0.0848) (**Figure 4S**). Maximum amplitude was modulated by grating movement, most strongly under open-loop (ON vs. OFF within OL, p < 0.0001; within CL, p = 0.0952) (**Figure 4T**). Together, these differences are consistent with reflexive OMR (during grating ON) depending on visuomotor contingency, whereas spontaneous swimming (during grating OFF) is less feedback dependent, and with functionally distinct motor-control pathways^31, 59^.

Finally, we validated the online model as a real-time detector. Running it on five WT AB fish at 5 dpf (N = 5 fish), we measured a mean bout-onset latency of 209.3 ± 2.9 ms (n = 291 bouts, mean ± s.e.m.), 211.3 ± 2.6 ms (n = 444 bouts), and 218.8 ± 3.9 ms (n = 175 bouts) for each of the sessions (**Figure S14A**), arising from the model’s input window (15 ms for three datapoints of future context), the feature-extraction pipeline (86.3 ms on average), and system-level processes such as shared-memory I/O. Although this latency remains to be optimized, 2-way ANOVA with Tukey’s *post-hoc* (α = 0.05) showed that only bout duration shifted relative to the Stytra-driven baseline, lengthening in all four conditions (CL-ON p = 0.0023; CL-OFF p = 0.0004; OL-ON p = 0.0120; OL-OFF p = 0.0006; Tukey), while frequency, interbout interval, tail beat frequency, and maximum amplitude were unaffected (all p > 0.32; 2-way ANOVA; **Figure S14B-U**). This duration increase is consistent with the bout lengthening reported under delayed visual feedback^20^.

### Analysis with Offline Model Reveals Trial and Strain Effects on Swim Bout Kinematics

Optical access to the brain is essential for recording neural activity during visuomotor behavior, and transparent mutants such as nacre (*mitfa^w^*^2^*^/w^*^2^, lacking melanophores) and casper (*mitfa^w^*^2^*^/w^*^2^; *roy^a^*^9^*^/a^*^9^, lacking melanophores and iridophores) are widely used for this purpose^60^. We compared head-fixed WT AB, nacre (NCR), and casper (CSP) larvae (N = 55, 31, and 26 fish) across four trials of alternating closed-loop/open-loop and grating ON/OFF conditions (**Figure 1F**). Per-fish, per-trial means were analyzed by 2-way ANOVA (strain × trial), followed by *post hoc* Tukey’s test (α = 0.05).

Bout frequency declined across trials in every condition (trial effect, all p < 0.0001), but this decline was evident in WT and NCR and not in CSP (**Figure 5A-D**). Under closed-loop grating movement (CL-ON), WT and NCR reduced bout frequency from Trial 1 to Trial 4 (WT p < 0.0001; NCR p = 0.0013), whereas CSP showed no significant change (p = 0.7807). By Trial 4, CSP therefore sustained a higher frequency than WT (p = 0.0284). The strain main effect and strain × trial interaction were not significant in any condition (e.g., CL-ON strain p = 0.3242, interaction p = 0.3893).

**Figure 5.**
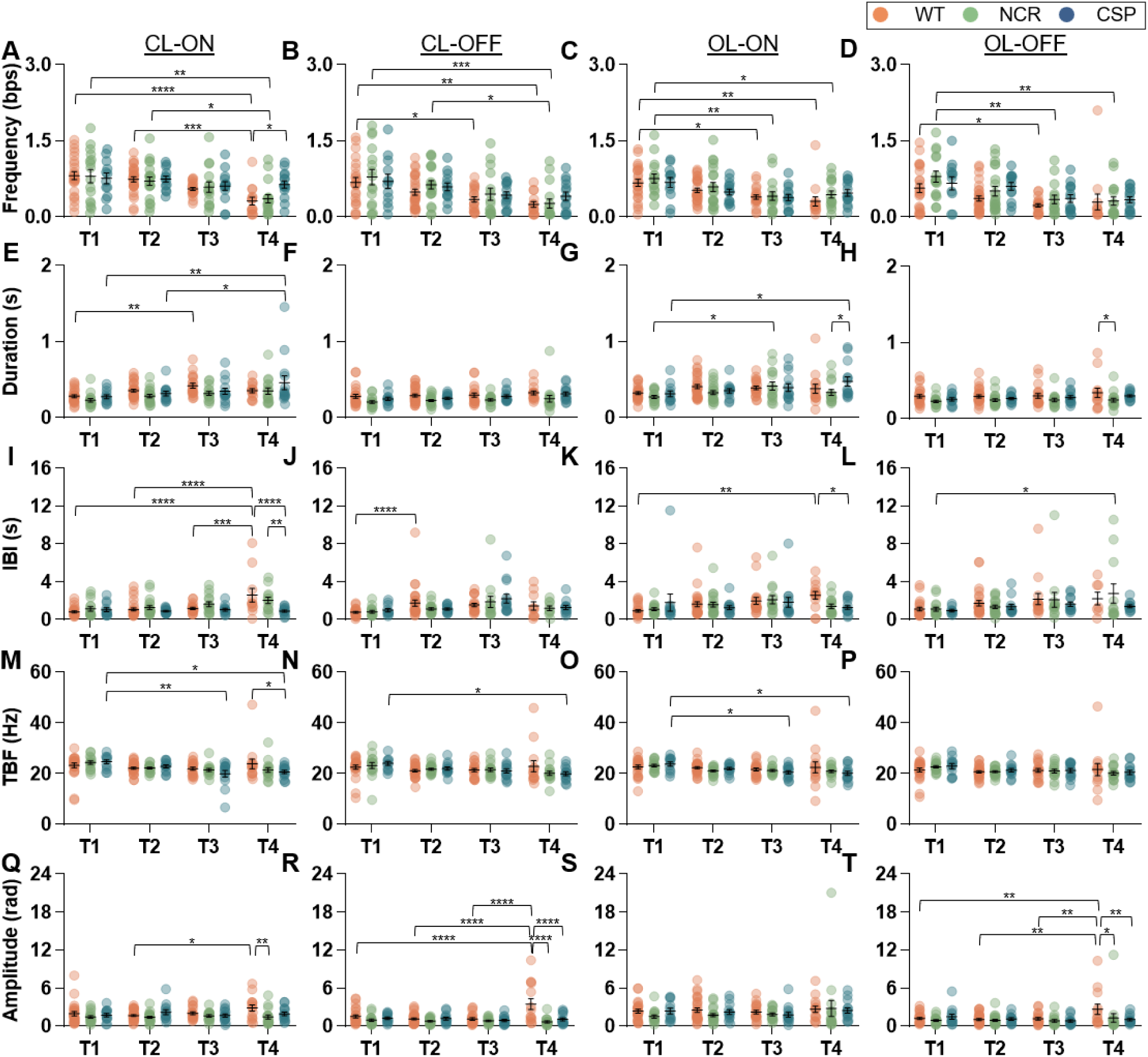
Trial number and genetic background have dissociable effects on swim-bout kinematics in WT AB, nacre, and casper larvae. Head-fixed larvae (N = 55 WT AB, 31 nacre [NCR], 26 casper [CSP], 6-7 dpf); four trials of three closed-loop + three open-loop (ON-OFF) cycles; closed-loop feedback driven by the Stytra threshold detector, with bout labels generated *post hoc* by the offline model. Per-fish, per-trial means; bars = mean ± SEM, points = single fish. Two-way ANOVA (strain × trial) with Tukey’s post-hoc; 95% CIs are of the pairwise mean difference. Panels are arranged by kinematic (rows) and condition CL-ON / CL-OFF / OL-ON / OL-OFF (columns). **(A-D) Bout frequency** declines across trials in every condition (trial main effect all p < 0.0001), a decline evident in WT and nacre but not casper. Under closed-loop grating movement **(A)**, WT and nacre reduce frequency from Trial 1 to Trial 4 (WT T1 > T4: mean difference 0.4987, 95% CI 0.2089-0.7885, p < 0.0001; nacre T1 > T4: mean difference 0.4417, 95% CI 0.1370-0.7464, p = 0.0013) whereas casper does not, so by Trial 4 casper sustains a higher frequency than WT (T4 WT < CSP: mean difference -0.3244, 95% CI -0.6213 to -0.02754, p = 0.0284). The strain main effect and strain × trial interaction are not significant in any frequency condition (CL-ON strain p = 0.3242, interaction p = 0.3893; CL-OFF strain p = 0.2376, interaction p = 0.9407; ; OL-ON strain p = 0.3025, interaction p = 0.8943; ; OL-OFF strain p = 0.0516, interaction p = 0.8500). Additional within-strain trial declines: WT T2 > T4 (**A**: mean difference 0.4265, 95% CI 0.1444-0.7086, p = 0.0007), nacre T2 > T4 (**A**: mean difference 0.3444, 95% CI 0.05532-0.6334, p = 0.0123); WT T1 > T3 (**B**: mean difference 0.3355, 95% CI 0.03550-0.6356, p = 0.0216) and T1 > T4 (**B**: mean difference 0.4359, 95% CI 0.1078-0.7640, p = 0.0039), nacre T1 > T4 (**B**: mean difference 0.5188, 95% CI 0.1766-0.8609, p = 0.0007) and T2 > T4 (**B**: mean difference 0.3683, 95% CI 0.04371-0.6930, p = 0.0191); WT T1 > T3 (**C**: mean difference 0.2788, 95% CI 0.04336-0.5143, p = 0.0130) and T1 > T4 (**C**: mean difference 0.3616, 95% CI 0.1037-0.6194, p = 0.0020), nacre T1 > T3 (**C**: mean difference 0.3451, 95% CI 0.06952-0.6207, p = 0.0075) and T1 > T4 (**C**: mean difference 0.3156, 95% CI 0.04453-0.5867, p = 0.0152); WT T1 > T3 (**D**: mean difference 0.3410, 95% CI 0.05624-0.6257, p = 0.0117), nacre T1 > T3 (**D**: mean difference 0.4504, 95% CI 0.1172-0.7836, p = 0.0032) and T1 > T4 (**D**: mean difference 0.4812, 95% CI 0.1534-0.8091, p = 0.0011). **(E-H) Bout duration** varies by trial (CL-ON p = 0.0001; OL-ON p = 0.0070) and by strain in three of four conditions (CL-ON p = 0.0297; CL-OFF p = 0.0003; OL-OFF p = 0.0010; OL-ON p = 0.1770, n.s.). Casper progressively lengthens its bouts across trials: CSP T1 < T4 (**E**: mean difference -0.1844, 95% CI -0.3274 to -0.04132, p = 0.0055), CSP T2 < T4 (**E**: mean difference -0.1435, 95% CI -0.2835 to -0.003422, p = 0.0424), CSP T1 < T4 (**G**: mean difference -0.1621, 95% CI -0.3157 to -0.008466, p = 0.0342); whereas WT and nacre change little (WT T1 < T3 in **E**: mean difference -0.1383, 95% CI -0.2500 to -0.02669, p = 0.0084; nacre T1 < T3 in **G**: mean difference -0.1416, 95% CI -0.2819 to -0.001249, p = 0.0471). Cross-strain differences emerge at Trial 4: NCR < CSP under OL-ON (**G**: mean difference -0.1436, 95% CI -0.2739 to -0.01342, p = 0.0266) and WT > NCR under OL-OFF (**H**: mean difference 0.09629, 95% CI 0.002587-0.1900, p = 0.0425). For CL-OFF **(F)** the strain main effect is significant (p = 0.0003) but no Tukey pair survives correction. **(I-L) Interbout interval** lengthens across trials in WT and nacre but not casper. **(I)** Under closed-loop grating movement, this produces a significant strain × trial interaction (p = 0.0115) and strain main effect (p = 0.0055): WT lengthens its interval from Trial 1 to Trial 4 (WT T1 < T4: mean difference -1.749, 95% CI -2.607 to -0.8915, p < 0.0001; T2 < T4: mean difference -1.499, 95% CI -2.337 to -0.6605, p < 0.0001; T3 < T4: mean difference -1.387, 95% CI -2.280 to -0.4945, p = 0.0005) while casper maintains a shorter interval than both WT (T4 WT > CSP: mean difference 1.670, 95% CI 0.7901-2.549, p < 0.0001) and nacre (T4 NCR > CSP: mean difference 1.128, 95% CI 0.2864-1.971, p = 0.0051). Casper also maintains a shorter interval than WT under OL-ON (**K**, T4 WT > CSP: mean difference 1.280, 95% CI 0.005285-2.554, p = 0.0488). Other within-strain calls: WT T1 < T2 (**J**: mean difference - 0.9651, 95% CI -1.816 to -0.1141, p = 0.0192); WT T1 < T4 (**K**: mean difference -1.630, 95% CI -2.890 to -0.3707, p = 0.0053); nacre T1 < T4 (**L**: mean difference -1.643, 95% CI -3.261 to -0.02534, p = 0.0450). Trial main effects: CL-OFF p = 0.0011, OL-OFF p = 0.0243; OL-ON n.s. (p = 0.0955). **(M-P) Tail-beat frequency** declines modestly across trials (CL-ON p = 0.0005; CL-OFF p = 0.0141; OL-ON p = 0.0050; OL-OFF p = 0.1082, n.s.) with no strain main effect in any condition (all p > 0.32). Significant pairwise: WT > CSP at CL-ON Trial 4 (**M**: mean difference 3.185, 95% CI 0.06892-6.302, p = 0.0439); within casper CL-ON T1 > T3 (**M**: mean difference 4.710, 95% CI 1.151-8.269, p = 0.0041) and T1 > T4 (**M**: mean difference 4.005, 95% CI 0.4457-7.564, p = 0.0205), CL-OFF T1 > T4 (**N**: mean difference 4.080, 95% CI 0.3718-7.787, p = 0.0247), OL-ON T1 > T3 (**O**: mean difference 3.265, 95% CI 0.05564-6.474, p = 0.0445) and T1 > T4 (**O**: mean difference 3.689, 95% CI 0.4794-6.898, p = 0.0171). **(Q-T)** Maximum amplitude shows a strain effect in three of four conditions (CL-ON p = 0.0039; CL-OFF p < 0.0001; OL-OFF p = 0.0412; OL-ON p = 0.3644, n.s.), with WT reaching higher amplitudes than both mutants at Trial 4 under CL-OFF (**R**: WT > NCR: mean difference 2.823, 95% CI 1.850-3.795, p < 0.0001; WT > CSP: mean difference 2.425, 95% CI 1.420-3.429, p < 0.0001) and under OL-OFF (**T**: WT > NCR: mean difference 1.361, 95% CI 0.2131-2.509, p = 0.0155; WT > CSP: mean difference 1.626, 95% CI 0.4414-2.810, p = 0.0040); WT > NCR also at CL-ON Trial 4 (**Q**: mean difference 1.406, 95% CI 0.3996-2.412, p = 0.0033), and within WT, CL-ON amplitude rose from Trial 2 to Trial 4 (**Q**, WT T2 < T4: mean difference -1.189, 95% CI -2.176 to -0.2020, p = 0.0111). The CL-OFF interaction is significant (**R**, p < 0.0001), driven by a WT-specific Trial 4 amplitude rise (WT T1 < T4: mean difference -1.959, 95% CI -2.948 to -0.9702; T2 < T4: mean difference -2.361, 95% CI -3.316 to -1.407; T3 < T4: mean difference - 2.359, 95% CI -3.387 to -1.332; all p < 0.0001).

Bout duration varied by trial (trial effect, CL-ON p = 0.0001; OL-ON p = 0.0070) and by strain in three of four conditions (strain effect, CL-ON p = 0.0297, CL-OFF p = 0.0003, OL-OFF p = 0.0010; n.s.: OL-ON p = 0.1770) (**Figure 5E-H**). CSP progressively lengthened its bouts across trials (CL-ON T1 vs. T4, p = 0.0055; T2 vs. T4, p = 0.0424; OL-ON T1 vs. T4, p = 0.0342), whereas WT and NCR changed little. NCR tended to produce the shortest bouts, reaching significance at Trial 4 in two conditions (WT > NCR under OL-OFF, p = 0.0425; CSP > NCR under OL-ON, p = 0.0266).

Interbout interval lengthened across trials in WT and NCR but not in CSP (**Figure 5I-L**). WT increased its IBI from Trial 1 to Trial 4 (CL-ON p < 0.0001; OL-ON p = 0.0053), and NCR did so under OL-OFF (p = 0.0450), whereas CSP’s IBI remained unchanged. Under closed-loop grating movement (CL-ON), this pattern produced a significant strain × trial interaction (interaction, p = 0.0115) and strain main effect (strain effect, p = 0.0055), with CSP maintaining a shorter IBI than both WT (p < 0.0001) and NCR (p = 0.0051) at Trial 4. CSP also showed a shorter IBI than WT under OL-ON at T4 (p = 0.0488).

Tail beat frequency declined modestly across trials (e.g., CL-ON p = 0.0005), with no strain main effect in any condition (strain effect, p > 0.32) (**Figure 5M-P**). Maximum amplitude showed a strain effect in three of four conditions (strain effect, CL-ON p = 0.0039, CL-OFF p < 0.0001, OL-OFF p = 0.0412; n.s.: OL-ON p = 0.3644), with WT reaching higher amplitudes than both mutants at Trial 4 (e.g., CL-OFF WT vs. NCR and WT vs. CSP, both p < 0.0001) (**Figure 5Q-T**).

Together, prior trial experience and genetic background had dissociable effects on bout kinematics. Trial effects modulated all five measured parameters, whereas strain effects were concentrated in bout duration and maximum amplitude. Casper did not show the late-trial decline in bout frequency and interbout interval observed in wildtype and nacre; instead, by Trial 4 it sustained bout frequency, maintained a shorter interbout interval, and produced longer bouts. This late-trial profile was therefore distinct from the other two lines, although whether it reflects the casper genetic background or other factors remains to be determined.

### The Choice of Bout Detector Alters the Biological Conclusions

Having adopted the best threshold method (Pow10S; **Figure S12A**) as a static reference, we asked whether the choice of detector changes the biological conclusions drawn from the same recordings. We re-ran the *adgrl3.1* KO and strain (WT/NCR/CSP) comparisons of **Figures 4** and **5** using the best threshold method in place of the offline model and compared the two detectors’ significance calls cell by cell (**Figures S15**; **Tables S9-S12**).

The two detectors agreed on the large majority of *post hoc* pairwise comparisons but returned different significance calls on 22: the offline model uniquely reached significance on 13 and the threshold method on 9. The divergence was kinematic-specific.

For the *adgrl3.1* positive control, the offline model uniquely detected two duration contrasts that the threshold method missed: the Trial 4 WT vs *adgrl3.1* KO difference under closed-loop (offline p = 0.0211, threshold p = 0.5051) and the OL-ON strain main effect (strain effect, offline p = 0.0016, threshold p = 0.2171), with no contrast detected by the threshold method (**Figure 4A-D** vs **Figure S15A-D; Table S9**).

For bout frequency, the offline model reported more significant differences (**Figure 5A-D** vs **Figure S15E-H**; **Table S10**): it uniquely detected five contrasts the threshold method missed, including the across-trial declines in WT and NCR (e.g., NCR CL-ON T2 vs T4, offline p = 0.0123, threshold p = 0.1240) and the cross-strain difference at Trial 4 (CL-ON WT vs CSP, offline p = 0.0284, threshold p = 0.0980), whereas the threshold method uniquely reached significance on only one (WT OL-OFF T1 vs T4, offline p = 0.1202, threshold p = 0.0434).

For bout duration the two detectors showed more divergence, each detecting four contrasts the other missed (**Figure 5E-H** vs **Figure S15I-L**; **Table S11**): the model uniquely reached significance on, for example, the CSP OL-ON T1 vs T4 lengthening (offline p = 0.0342, threshold p = 0.9477), while the threshold method uniquely reached significance on the Trial 1 and 2 WT vs NCR under closed-loop grating pause (offline p = 0.0612 and 0.0838, threshold p = 0.0074 and 0.0110).

For interbout interval (IBI), the threshold method reached significance on more contrasts than the model (four vs two) (**Figure 5I-L** vs **Figure S15M-P**; **Table S12**). The threshold method uniquely reached significance on a within-casper closed-loop progression that the model did not (e.g., CSP CL-OFF T1 vs T3, offline p = 0.0507, threshold p = 0.0092), whereas the model uniquely reached significance on the closed-loop strain × trial interaction underlying the CSP IBI pattern described above (interaction, offline p = 0.0115, threshold p = 0.1626) and on the NCR OL-OFF T1 vs T4 lengthening (offline p = 0.0450, threshold p > 0.9999).

Because ground-truth labels are not available for every session in these larger cohorts, we do not claim that either detector is uniformly correct, nor do we interpret any individual divergence as a true or false call. The comparison instead shows that the choice of bout detector leads to different statistical analysis results from the same dataset, in a kinematic-specific pattern: for bout frequency the offline model reached significance on more contrasts (five detected only by the model vs one only by the threshold method), for bout duration the two detectors diverged symmetrically (each reached significance on four contrasts the other missed), and for interbout interval the threshold method reached significance on more contrasts (four vs two). A threshold pipeline’s conclusions further depend on the threshold, window, and tuning chosen, whereas the threshold-free model returns a single call per dataset without per-experiment tuning.

## Discussion

We developed two threshold-free bout detectors for larval zebrafish: a high-accuracy offline model for *post hoc* analysis and a lightweight online model for real-time closed-loop use. Both operate on tail-curvature dynamics rather than a manually tuned threshold. The offline model’s advantage is most evident in low-amplitude sessions, where genuine bouts and tracking noise overlap in magnitude, and any single threshold must trade false negatives against false positives. By leveraging temporal context rather than instantaneous amplitude, the model reduces both error types simultaneously. Because it requires no per-experiment tuning, the models also yield a single, reproducible result across datasets, removing a user-dependent source of variability from downstream biological analysis.

Applied to our head-fixed, closed-loop preparation, the model shows that head fixation preserves the primary features of free-swimming locomotion: bout frequency, duration, interbout interval, and tail-beat frequency did not differ from freely swimming larvae. The exception was an increase in maximum bout amplitude under restraint, likely because agarose prevents forward translation, so propulsive effort is expressed as greater tail bending rather than body displacement; this hypothesis, however, requires dedicated biomechanical testing. Overall, the preparation remains suitable for paradigms requiring head fixation.

Across repeated trials, locomotor output increased between the first two trials, potentially due to initial acclimation to the arena^61, 62^, and then declined steadily. Two timescales were evident. The abrupt within-trial drop at open-loop onset – when concordant visual feedback is removed – is consistent with a sensory-prediction signal detecting mismatches between expected and actual optic flow^33^. The slower across-trial decline is less easily attributed to a single mechanism and is compatible with non-associative habituation, accumulating fatigue, or stress adaptation, which cannot be disentangled in the present design.

Genetic background shapes zebrafish phenotypes^2, 63^, including drug responses^64–66^, anxiety^67^, social interaction^68, 69^, escape^8, 70^, and navigation^71^, yet its influence on the optomotor response has remained largely uncharacterized. A prior pigmentation-mutant comparison being limited to bout frequency^60^. Across wildtype, nacre, and casper larvae, trial experience and strain background acted differently: trial experience broadly modulated bout kinematics, whereas strain effects were concentrated in bout duration and maximum amplitude, with a strain-by-trial interaction in closed-loop interbout interval. Notably, casper did not exhibit the late-trial decline in frequency and interbout interval observed in wildtype and nacre. Transparent imaging strains are therefore not behaviorally interchangeable with wildtype AB fish – at least within our OMR head-fixed assay – and behavioral conclusions should be validated before extrapolating across genetic backgrounds.

In real-time use, the online model operates within the closed loop but currently incurs ∼200 ms of bout-onset latency, driven primarily by feature extraction and interprocess communication. Consistent with prior reports that delayed visual feedback lengthens bouts^20^, this latency selectively increased bout duration without affecting other kinematic parameters. Network quantization and integration of tracking and inference into a unified software pipeline, eliminating interprocess overhead, should reduce latency toward true real-time performance.

Two limitations define the present scope. First, although all larvae were tested at 5-7 dpf, detection accuracy is governed more strongly by bout amplitude than by age: model performance was most challenged by the smallest bouts near the tracking-noise floor, so transfer across stages, strains, or recording systems will depend on overlap in tail_sum amplitude with the training data. Second, the free-swimming pipeline does not yet co-register the 35 mm arena boundary, preventing automatic exclusion of wall-proximal bouts. This limitation is expected to be resolved in the integrated tracking software under development.

Finally, broader cross-laboratory and cross-genotype generalization will require larger labeled datasets than currently available. Here, the *adgrl3.1* knockout served as a positive control rather than a phenotyping target. To enable community-driven extension, we release both the labeled dataset and source code under an MIT license. Because the detector segments bouts directly from continuous tail kinematics without a threshold, it can function upstream of probabilistic and deep-learning classifiers of bout subtype, such as Megabouts^24^. Pairing threshold-free segmentation with downstream classification is a clear step toward fully automated, reproducible analysis of larval swimming.

## Methods

### Larval Zebrafish

All experimental procedures were conducted in accordance with National Institutes of Health (NIH) guidelines and were approved by the Johns Hopkins University Animal Care and Use Committee (ACUC, approval number can be requested through the corresponding author). Larval zebrafish (Danio rerio) of wild-type (WT AB), nacre (*mitfa^w^*^2^*^/w^*^2^), and casper (*mitfa^w^*^2^*^/w^*^2^; *roy^a^*^9^*^/a^*^9^) genetic backgrounds were raised and maintained in-house. Adult fish were maintained at 28.5 °C on a 14:10 hour light-dark cycle; embryos and larvae were raised in E3 medium at 28.5 °C. Fish were tested at 5 days post-fertilization (dpf) for experiments using the online bout-detection model to detect bouts in real time, 12 dpf for tricaine-treated negative control, and at 6-7 dpf for all other behavioral experiments. Sex is not a relevant variable at the larval stage, as sexual differentiation does not occur until 3-4 weeks of age. At the end of each experiment, larvae were euthanized by immersion in dilute bleach (sodium hypochlorite) on ice.

### Control and Experimental Groups

Eleven WT AB larvae (N = 11 fish) were used to establish baseline free-swimming behavior. Fifty-five WT AB (N = 55 fish), thirty-one nacre (N = 31 fish), and twenty-six casper (N = 26 fish) larvae were used for the head-fixed strain comparison. The head-fixed WT AB group served a dual purpose: (1) as the experimental group for comparison against the free-swimming condition to assess the effects of mounting, and (2) as the control group for evaluating kinematic differences relative to the nacre and casper strains. As a positive control for behavioral sensitivity, eleven *adgrl3.1* G0 knockout (KO) larvae (AB background; N = 11 fish; hereafter *adgrl3.1* KO), assayed head-fixed at 7 dpf, were compared against the WT AB group (**Figure 4A-H; Figure S13**). As a negative control, five larvae (N = 5 fish, four 12 dpf and one 7 dpf, WT AB) were recorded under tricaine anesthesia (**Figure 4I**). A separate cohort of five WT AB larvae (N = 5 fish, 5 dpf) was recorded under real-time online-model control (**Figure S14**). The model training and test recordings (8 and 6 head-fixed WT AB larvae; see ***Model Training and Validation***) partially overlap with the head-fixed WT AB cohort. Per-figure cohort assignments, ages, and recording conditions are given as dataset tags in the corresponding figure captions.

### Adgrl3.1 Knockout

Fish for *adgrl3.1* KO experiments were generated following an established G0 KO method^72^. Briefly, four distinct guide cut sites were selected on the basis of combined on-/off-target criteria derived from the CRISPOR^73^ and CHOPCHOP^74^ online tools. Oligonucleotides (Integrated DNA Technologies, USA) were used to generate sgRNA templates by annealing and elongation of a primer containing a T7 promoter and guide sequence together with tracrRNA for the standard sgRNA scaffold (Phusion High-Fidelity DNA Polymerase, ThermoFisher, USA). The guide sequences used are listed below:

**Guide 1:**

TAATACGACTCACTATAGGCTGATGCCTACAAGATTAGTTTTAGAGCTAGAAATAGC

**Guide 2:**

TAATACGACTCACTATAGGTCTTATTGGATGACTGTAGTTTTAGAGCTAGAAATAGC

**Guide 3:**

TAATACGACTCACTATAGGCCTCCACTCTGAGTGTGTGTTTTAGAGCTAGAAATAGC

**Guide 4:**

TAATACGACTCACTATAGGAGTCGTGGAGCTGCGTACGTTTTAGAGCTAGAAATAGC

Templates were in vitro transcribed overnight (HiScribe T7 High Yield RNA Synthesis Kit, New England Biolabs, USA) and purified (RNA Clean & Concentrator, Zymo, USA). A working solution was prepared by combining all four sgRNAs with EnGen Spy Cas9 NLS (New England Biolabs, USA) and diluting with RNase-free water to a final concentration of 2.5 μM Cas9 and 1 μg/μL total sgRNA. The mix was incubated at 37 °C for 10 min to form Cas9-ribonucleoprotein complexes. Approximately 1 nL was microinjected into the cytoplasm of early one-cell-stage embryos, which were moved to a 27 °C incubator immediately following injection.

### Tricaine Treatment

As a negative control in which locomotor activity is suppressed, five larvae (N = 5 fish; four at 12 dpf, one at 7 dpf; WT AB) were head-fixed and recorded under tricaine (MS-222) anesthesia (**Figure 4I**). A 1× tricaine (MS-222) solution in E3 medium was prepared per the recipe of Schoen et al^75^. After tricaine application, larvae were held for the same 5-10 min interval as the standard pre-assay acclimation (see ***Zebrafish Larva Mounting Protocol for Head-Fixation***) before the behavioral protocol began, so that treated and untreated conditions differed only in the presence of anesthetic.

### High-Framerate Visual Stimulus Generation

To achieve a stimulus refresh rate exceeding the 60 Hz limit of Stytra^34^, we set up inter-process communication pairing Stytra (Python 3.8) with PsychoPy^38^ (Python 3.10+). Communication was managed using the multiprocessing.shared_memory module in Python. The primary Stytra process allocated shared-memory buffers to relay commands and real-time data to the concurrent PsychoPy process, enabling instantaneous updates to the visual stimulus (**Figure S2**, **Table S1**). The Stytra-PsychoPy workflow was used for both free-swimming and head-fixed OMR protocols, and the Stytra bout detector was used for real-time visual feedback for all head-fixed fish except in **Figure S14**.

### Free-Swimming Setup

Experiments on free-swimming larvae were conducted in the arena depicted in **Figure 1A**. The arena was illuminated from below by a custom infrared (IR) LED array (Waveform Lighting, USA). A high-speed camera (Teledyne, USA) positioned above the arena captured dorsal-view images of the fish at 200 Hz (1440 × 1080 pixel resolution). A varifocal lens (Computar, USA) and a long-pass filter (LPF_1, λp ≥ 830 nm, Schott, Germany) were mounted to the camera. The camera field of view was adjusted to encompass the entire 35 mm Petri dish (FABLAB, USA) containing the larva. The 35 mm dish (approximately 10× the larval body length) lies within the range of arenas used for larval zebrafish locomotor assays^2, 8, 31, 60, 68, 71, 76^. Visual stimuli, generated by PsychoPy, were displayed using a projector (KODAK, USA) onto a cold mirror (DM, Techspec, USA) and then reflected to a silicone diffusion pad beneath the Petri dish. A long-pass filter (LPF_2, λp ≥ 590 nm, Thorlabs, USA) in the projection path created a high-contrast red-on-black stimulus, known to effectively elicit visuomotor responses^39, 40^.

### Free-Swimming Optomotor Response (OMR) Protocol

A single larval zebrafish (6-7 dpf) was placed in a 35 mm Petri dish with E3 medium at 29 °C. After a 5-10 minute acclimation period with a static grating, a 480 s protocol was initiated (**Figure 1E**). The protocol consisted of six trials, each comprising two grating movement-pause cycles with opposite movement directions. Each cycle comprised 20 s of grating movement (spatial period ≈ 4 mm, speed ≈ 4 mm/s) followed by a 20 s static pause, and the two cycles within a trial used opposite directions (e.g., 90° ± 15° then 270° ± 15°) to prevent the fish from dwelling at the dish edge.

### Head-Fixed Setup

For experiments with head-fixed fish, a compact version of the behavioral setup was designed for integration with microscopes^7^, hereafter referred to as the “closed-loop setup” (**Figure 1B**, **Figure S1**). Tail motion was tracked using a folded IR imaging path. An IR LED array (Waveform Lighting, USA) was positioned next to the microscope objective to illuminate the tail of the fish. The IR light first passed through the cold mirror (DM, Techspec, USA) and was then redirected by a plane mirror (PM, Thorlabs, USA). Images were captured at 200 Hz by a high-speed camera (Teledyne, USA) through a custom 4f optical system (a camera lens from Kowa, Japan, focused at infinity, followed by a 175 mm biconvex lens L1 from Thorlabs, USA) and a long-pass filter (LPF_1, λp ≥ 830 nm, Schott, Germany). To accommodate the different working distances of the 10× and 20× OPM objectives, L1 was mounted on a slider for manual focus adjustment. As in the free-swimming setup, a long-pass filter (LPF_2, λp ≥ 590 nm, Thorlabs, USA) was used to create a high-contrast red-on-black visual stimulus, presented by reflection from a projector (Sony, Japan) via the cold mirror (DM) onto a projection screen. The projection screen was constructed from weighing paper (Fisherbrand, USA) with a rectangular aperture, ensuring the tail of the fish remained visible to the camera while allowing the visual stimulus to be presented on the surrounding surface. The position of the aperture was adjusted with a slider on the 3D-printed dish holder.

### Zebrafish Larva Mounting Protocol for Head-Fixation

To ensure consistent and minimally invasive head-fixation for behavioral assays, a mounting protocol was adapted from established methods^41–44^. Larval zebrafish (6-7 dpf) were selected for vigor, identified by their active escape response to gentle suction with a transfer pipette. A selected larva was transferred into 2% (w/v) low-melting-point agarose (LMPA, Fisher Scientific, USA) held at 42 °C. Approximately 35 μL of agarose containing the larva was pipetted into the center of a Petri dish (FABLAB, USA). Before solidification, the larva was quickly oriented into a dorsal- up position by gently tapping the Petri dish or using a soft, fine-tipped tool (e.g., a fire-polished pipette tip). The agarose was allowed to solidify for approximately one minute (**Figure S3A**).

Next, a custom 3D-printed tool was used to cut the agarose and expose the tail and eyes (**Figure S3I**). Both the tail and the eyes were freed from agarose: the tail to allow unrestrained tail kinematics, and the eyes to reduce mechanical restraint and to keep the preparation compatible with optokinetic response (OKR)-based studies, in which eye movements must remain unobstructed (eye motion during our OMR assay was also observed in **Figure S3L-M**). To free the tail, four symmetrical, angled cuts (∼45° distally) were made: two at the mid-caudal point of the body and two originating near the pectoral fins (**Figure S3B**). The ideal agarose texture allows for a clean incision without exudation of liquid; if the cut is not clean, additional solidification time is recommended. Following the incision lines, the blade was inserted vertically and used to gently peel the agarose block away from the body, freeing the entire caudal region of the fish (**Figures S3C-E**). A similar procedure was used to expose the eyes: angled cuts were made proximal to the eyes (**Figure S3F**), and the blade was used to carefully lift and remove the resulting agarose pieces, ensuring an unobstructed field of view (**Figures S3G-H**). All cuts were made approximately 1 mm away from the fish; the rigidity of LMPA allows complete removal of the excess piece along the cuts.

Immediately after agarose removal, a small drop (∼35 μL) of E3 medium was applied to the fish to prevent drying. Before the behavioral assay, the Petri dish was filled with approximately 2 mL of warm (29 °C) E3 medium, added gradually to avoid dislodging the larva from the agarose. Mounted larvae were then acclimated for 5-10 minutes under a static grating, and recording began only once intermittent spontaneous bouts were visible and successfully tracked in the Stytra preview, confirming behavioral readiness.

### Fixed-Head Optomotor Response (OMR) Protocol

Head-fixed fish (6-7 dpf) were acclimated for 5-10 minutes under a static grating (see ***Zebrafish Larva Mounting Protocol for Head-Fixation***) before starting a 960 s session (**Figure 1F**). The session consisted of four identical trials, each containing three closed-loop (CL) cycles followed by three open-loop (OL) cycles. Each cycle included a 20 s stimulus-movement period (grating ON) and a 20 s stationary period (grating OFF). During grating ON, the grating moved tail-to-head to drive forward optomotor swimming.

- Closed-Loop (CL): the grating speed (spatial period = 4 mm, base velocity ≈ 4 mm/s during grating ON and 0 during grating OFF) was dynamically updated based on the fish tail movements to create a virtual swimming environment, driven by real-time bout detection (described below).
- Open-Loop (OL): the grating moved forward at a constant velocity of ≈ 4 mm/s during the ON phase and was stationary during the OFF phase.

### Real-Time Tail Tracking and the Tail_sum Input Signal

Tail kinematics were tracked in Stytra^34^, which represents the tail as a series of segment angles (*θ*_0_ to *θ*_5_; **Figure S5**). Stytra was configured with *n_segments* = 18 (segments fit during skeletonization) and *n_output_segments* = 6 (output tail angles), following prior larval-zebrafish work^77^; these two parameters were fixed for all recordings. The remaining image-preprocessing and tracking parameters (gain, binarization threshold, region of interest, and body and tail-start orientation) were adjusted at the start of each session to maintain tracking quality across the varied imaging conditions of the multi-month data acquisition, and the transparent mutant showed equivalent tracking success as to the wildtype (**Figure S4**). Tracking output and the corresponding tail_sum trace were inspected against the raw video qualitatively for every session before acceptance into the dataset.

The model input is based on Stytra’s centroid-based *tail_sum* (*θ*), defined as the sum of the two distal segment angles minus the sum of the two proximal segment angles:

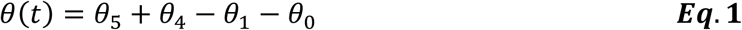

A frame-to-frame temporal difference was applied to this signal, which we refer to as tail_sum in this work for consistency throughout:

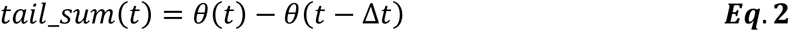

to suppress slowly varying components, such as low-frequency optic-table swing and illumination or sensor-baseline drift, which contribute equally to consecutive frames and cancel in the difference while retaining frame-to-frame tail motion.

For closed-loop feedback, the signal was fed in real time to Stytra’s threshold-based detector (to generate the datasets) or to the online model (for real-time validation). On detection, a signal was sent via shared memory to PsychoPy, which updated grating velocity by a custom function:

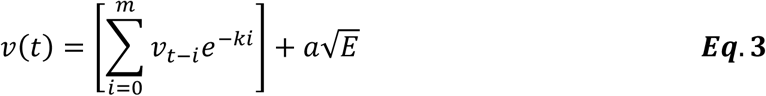

estimating the fish’s intended forward motion. The function integrates previous velocities to model inertia (*m* = 4, *k* = 1.0366) and adds thrust proportional to the instantaneous tail-beat energy *E* (scaling factor *a* = 0.3368), where *E* is the power of the 0.25*fs*-0.5*fs* frequency band of tail_sum from an 8-point short-time Fourier transform. These parameters were derived from free-swimming data during pilot run. The Stytra interface and the PsychoPy stimulus window were screen-recorded in parallel at 60 Hz (OBS Studio, Lain Bailey, USA) for post-hoc annotation (**Figure S5**).

### Manual Annotation of Bouts

Ground-truth labels were generated by a two-stage procedure using custom MATLAB scripts (MathWorks, R2024b). In the first stage, swim bouts were coarsely annotated frame by frame from the synchronized video of each head-fixed session in a graphical interface (*videoAnalysisTool*; **Figure S5A**); a bout was marked when the distal tail bent laterally relative to the stationary proximal tail. In the second stage, each bout’s onset and offset were refined from quantitative signal changes in a second tool (*boutExtraction*; **Figure S5B**), which displayed the raw tail_sum trace together with its moving sum of squares and 20-40 Hz bandpass filtered tail_sum. The first deflection point in the BPF signal corresponding to the labeled bout was defined as the bout onset, and the last deflection point before the raw signal returned to baseline (plateau) was defined as the bout offset. No amplitude or duration cutoff was applied, so that low-amplitude bouts were not excluded by construction. Candidate events rejected in the second stage were discarded rather than relabeled. Each timepoint was labeled bout (1) or no-bout (0). The training set comprised 8 head- fixed sessions (N = 8 fish, 6-7 dpf WT AB; 3743 bouts) and the held-out test set 6 sessions (N = 6 fish; 2948 bouts). One annotator provided label for training and testing set. Three annotators provided additional labels for inter-rater analysis. All annotators were blinded to the H/L amplitude classification of the sessions, as the classification was not established during the labeling process. Per-session composition and inter- and intra-rater agreement (Cohen’s κ and percent agreement) are reported in **Table S2**. The annotation training slide is included in the project repository.

### Feature Extraction and Model Architecture

Two models were developed: a high-accuracy offline model and a low-latency online model. Missing values in the tail_sum trace were imputed by quadratic spline interpolation and zero-padded to the length of the prediction sequence. A feature matrix was built with a sliding window (50 points offline, 30 points online), the offline model predicting the label of the window’s 31st point (the 27th online). Nine features were computed over a 3-point rolling window within the offline window (**Table S3**): seven time-series statistics (the rolling mean, maximum, minimum, sum of squared values, mean of absolute value, first derivative, and gradient of squared values)^34, 47–50^, the raw Stytra tracking signal, and the 20-40 Hz (4th-order Butterworth) band-pass filtered tail_sum power, whose passband targets the larval tail-beat frequency band^51^. The online model used a six-feature subset with the highest permutation feature importance (PFI, **Table S4**), and its computation is further accelerated with a just-in-time (JIT) compiler^78^. Candidate features were restricted to those computable in O(N) per window for real-time use; the final feature sets were selected from a larger initial pool by permutation feature importance (**Table S5**).

The feature matrix passes through three 1D convolutional blocks (filters 64, 128, 256; kernel widths 9, 3, 1; ReLU), each followed by max-pooling (×2) and dropout (0.2) (**Figure 2C**). In the offline model the convolutional output feeds a 128-unit bidirectional LSTM (BiLSTM) that integrates the full window in both directions; the BiLSTM is omitted in the online model to reduce inference latency. A final dense layer with sigmoid activation gives a per-timepoint probability, with values above 0.5 classified as bout. Omitting the BiLSTM raises the real-time tracking-and-detection rate from approximately 50 Hz (offline) to approximately 150 Hz (online).

Model parameters were fixed to match the previous studies^53, 54^ and verified on measured time scale of larval swimming. At 200 Hz, mean labeled bout durations were ≈ 280-420 ms (≈ 56-84 frames) and tail-beat frequency ≈ 21-24 Hz (≈ 8-9 frames per cycle), so the 50-point (250 ms) window spans roughly one bout and 5-7 tail-beat cycles (**Figure 4**). The convolutional kernels (9, 3, 1) capture structure at progressively coarser scales, and the BiLSTM aggregates the full window to distinguish sustained low-amplitude oscillations from transient noise. Parameter counts were 837,000 (offline) and 278,000 (online). The simpler baseline variants used for the architecture ablation are described in ***Simpler Machine-Learning Baselines*** (**Figure 3J**).

### Model Training and Validation

Models were trained with binary cross-entropy loss and the Adam optimizer (learning rate 1×10⁻⁴) for up to 100 epochs (batch size 64), with early stopping on validation F1 (*patience* 5) and a learning-rate scheduler. Data were split at the animal level: N = 8 head-fixed WT AB fish for training and N = 6 held-out head-fixed WT AB fish for testing, with no fish in both; both models were trained and validated on head-fixed recordings only. Model selection used 8-fold cross-validation over segmented windows within the training set (*shuffle* = True, *random_state* = 42). All reported performance is computed on the held-out test set. The simpler-baseline ablations (**Figure 3J**) used animal-level GroupKFold (n_splits = 8, one fish per fold) on the same partitions, whereas the logistic-regression baseline was fit once on the full standardized training set. All were evaluated on the same held-out test fish. Bout labels represented 14.2% of training frames and 16.1% of test frames. No class weighting or resampling was applied, and model selection used the F1 score rather than accuracy under this imbalance^79, 80^.

### Simpler Machine-Learning Baselines

Five simpler baselines were trained to isolate which components of the offline architecture contribute to detection. All used the same windowed input and labels as the offline model and were evaluated per fish on the same N = 6 held-out test fish (**Figure 3J**). The neural baselines used animal-level GroupKFold over the training fish (leave-one-fish-out), with the best fold selected by validation F1; the logistic regression was fit directly on the training set.

- Logistic regression (LR): an L2-regularized logistic regression (scikit-learn, L-BFGS solver) on the standardized, flattened 50-point × 9-feature window (450 inputs; 451 parameters).
- CNN: one 1D convolutional layer (64 filters, kernel width 9, ReLU, same padding) followed by dropout (0.2), max-pooling, and a sigmoid output (6,849 parameters), removing the stacked convolutional depth and the BiLSTM of the offline model.
- MLP on aggregated features (MLP-1): a three-layer perceptron (256, 128, 64 units; ReLU; dropout 0.2) on a 45-dimensional per-window summary - the mean, standard deviation, minimum, maximum, and instantaneous value of each of the nine features (52,993 parameters) - removing fine-grained temporal structure.
- MLP on label-time features (MLP-2): a three-layer perceptron (64, 32, 16 units; ReLU; dropout 0.2) on the nine features at the labeled timepoint only (3,265 parameters), removing all temporal context.
- Offline-2: the full offline architecture (three convolutional blocks and the BiLSTM) applied to the six raw tracked segment angles in place of the nine tail_sum-derived features, substituting the input derived from a single summarizing tail_sum signal of the entire tail curvature.

The neural baselines were trained with the Adam optimizer (learning rate 1×10⁻⁴) and binary cross-entropy loss for up to 100 epochs (batch size 64), with early stopping on the validation F1 and a learning-rate scheduler; the logistic-regression and MLP inputs were standardized. All models classified each timepoint at a 0.5 probability threshold.

### Inference and Threshold-Based Benchmark

For inference on new recordings, the same pipeline (interpolation, padding, feature extraction) was applied. Free-swimming recordings were split into contiguous tracked segments processed independently, with predictions during tracking failure set to not-a-number (NaN).

Each recording session (one head-fixed fish with one continuous tail_sum trace) was classified as high amplitude if more than half of its annotated bouts exceeded a peak-to-peak tail_sum amplitude of 1.0 rad, and low amplitude otherwise. The per-session Pearson correlation between bout peak-to-peak amplitude and time-within-session was computed to check for within-session amplitude drift (**Table S7**).

Detection was benchmarked against the threshold detector built into Stytra, which flags a bout when the rolling standard deviation of tail_sum over a 10-point window exceeds a fixed threshold (default 0.050). The benchmark comprised 32 additional threshold variants spanning four statistic families (rolling standard deviation, rolling mean, rolling median, and the moving average of the 20-40 Hz band-pass filtered tail_sum power)^51, 81, 82^, four window lengths (10, 50, 70, 100 points), and global or per-session calibration. Variants were tuned on the training set and evaluated on the low-amplitude test sessions; per-session adaptive calibration was performed directly on the testing set (**Figure S12A**). A per-session calibrated variant reached higher F1 on the low-amplitude subset but requires session-level labels and was not adopted as the reference. The F1-maximizing variant with a global threshold – a 10-point moving average of the 20-40 Hz band-pass filtered tail_sum power with a single global threshold – is referred to as the best threshold method (Pow10S) and was used as the static reference detector for all downstream biological comparisons. Generalization was assessed by evaluating the offline model and the Stytra detector on three labeled free-swimming sessions (N = 3 fish; **Table S8**). Benchmarking was performed without the post-processing steps below.

### Detection Post-Processing and Kinematic Parameter Extraction

For biological analysis, bouts were detected from the raw tail_sum trace with the offline model and then post-processed: segments shorter than 75 ms were discarded as noise, and segments longer than 2.0 s were split at points of inactivity or discarded^19^. The following kinematic parameters were computed from the curated bouts:

- Bout frequency (bout per second, bps): the number of bouts initiated within a condition. For free-swimming recordings, frequency was normalized by the total successfully tracked duration in the condition.
- Bout duration: the time (s) from the start to the end of a bout, which had to start and end within the same condition (e.g., entirely within a grating-ON period).
- Interbout interval (IBI, s): the time from the end of one bout to the start of the next, with both bouts within the same condition.
- Tail-beat frequency (TBF, Hz): the number of zero-crossings per unit time of the normalized tail_sum trace during the bout.
- Maximum amplitude (rad): the peak-to-peak tail_sum (maximum - minimum) within a bout.

### Statistics and Reproducibility

Statistical analyses were performed in Prism (v10.6.1, GraphPad, USA) at a significance level of α = 0.05. Individual fish were treated as biological replicate, and the sample size N reported in figures and text refers to the number of fish. Kinematic parameters were averaged per fish within each condition, and these per-fish means were used for all comparisons. Data are mean ± s.e.m. unless otherwise noted, and 95% confidence intervals for the difference of mean accompany each pairwise comparison reported in the captions of **Figures 4**-**5**, and **S13-14**.

Two-factor designs used an ordinary 2-way ANOVA with a full interaction model. Post-hoc pairwise comparisons used the Prism default uncorrected Fisher’s LSD test for 2×2 designs (**Figure 4K-T**) and Tukey’s HSD test for designs with three or more rows or columns (**Figure 4A-H**, **Figure 5**, **Figures S13-S15**), both from the same 2-way ANOVA in Prism.

- ***Sample size***. Sample sizes were not set by an a priori power analysis. Cohort sizes followed the range typical of larval-zebrafish kinematic studies (approximately 3 to 74 fish per group^2, 8, 31, 56, 60, 67–70, 76^. A post-hoc sensitivity power analysis (α = 0.05, 1 - β = 0.80; Python 3.11, *statsmodels*) gave the minimum detectable effect size (MDES) for each comparison family (**Table S13**): Cohen’s f = 0.2974 to 0.5223, partial η² = 0.0812to 0.2144 for the 2-way ANOVA main and interaction effects across the five test families, with Fisher’s LSD pairwise Cohen’s d = 0.5391 to 1.2560.
- ***Inclusion and exclusion.*** Larvae were screened for vigor (escape response to gentle suction) before all sessions (free swimming and head-fixed); no other *a priori* inclusion criterion was applied. Recordings with persistent tracking failure were excluded. During post-processing, segments shorter than 75 ms were discarded and segments longer than 2.0 s were split or discarded.
- ***Randomization and blinding.*** Genotype and treatment groups were defined by design and were not randomized; within each genotype, larvae were drawn arbitrarily from the housing population then screened for vigor. Closed-loop and open-loop blocks were presented in a fixed order within each head-fixed session, and free-swimming grating directions alternated between cycles yet with randomized orientation around the target direction. Annotators were blinded to genotype except the first author; recording condition (grating ON/OFF, closed- versus open-loop) was not visible in the recordings.

## Data Availability

Data contrasting the figures are available at Supplementary Data. Datasets, labels, and model predictions are available at https://github.com/yibiophotonics/Threshold-free-bout-detection.

## Code Availability

Code for data acquisition and analysis are available at https://github.com/yibiophotonics/Threshold-free-bout-detection

## Author Contributions

J.Y. and H.W. conceptualized the study; H.W. and Y.Z. established the free-swimming behavioral arenas and revised the tracking code; Y.Z. developed algorithm to transform tracking data to fish velocity; H.W. designed and implemented the fixed-head behavioral setup, experimental protocols, and stimuli display; G.K. bred and raised the larval zebrafish; H.W. and G.K. jointly established the fish mounting protocol; H.W. performed the *in vivo* larval zebrafish experiments; H.W. designed, trained, and validated the deep learning models; H.W., G.K., and J.Y. performed data analysis; H.W. and J.Y. prepared the figures and wrote the manuscript; all authors contributed to manuscript revision; J.Y. and J.S.M. coordinated the collaboration, and J.Y. provided overall supervision for the study.

## Acknowledgement

This manuscript was supported in part by the National Institutes of Health (NIH, EBEB034272; AG072305; NS041435) and subject to the NIH Public Access Policy. NIH has been given the right to make this manuscript publicly available in PubMed. We also acknowledge the Boons Pickens Endowment Fund, and an unrestricted grant to the Wilmer Eye Institute from Research to Prevent Blindness. GK received support from the Johns Hopkins Cross-Disciplinary Graduate Program in Biomedical Sciences. JSM received support from the Helen Larson and Charles Glenn Grover endowment funds.

