## Supplementary Material for "Threshold-Free Neural Network Models for Swim Bout Detection of Larval Zebrafish"

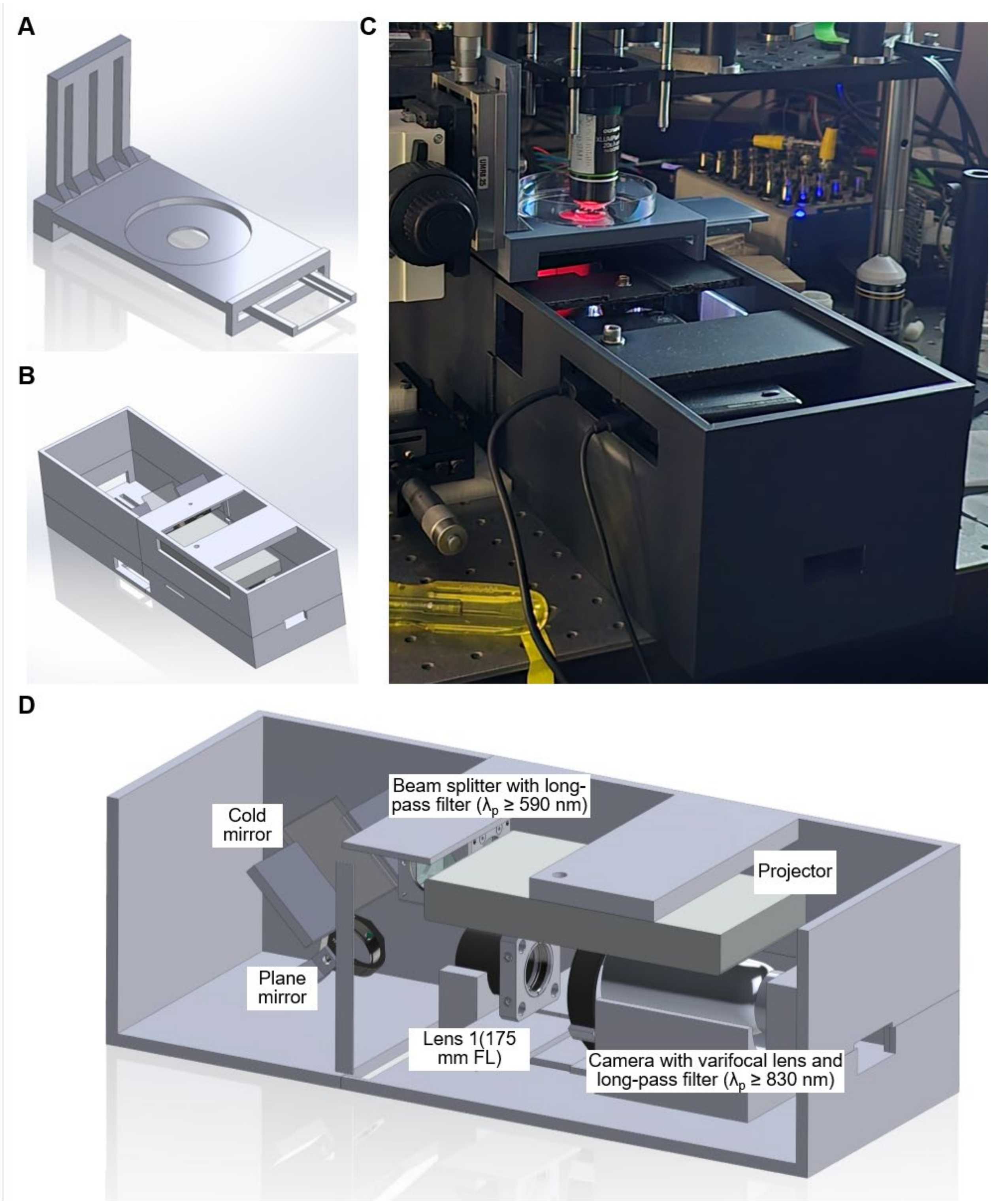


**Figure S1. Integration of the head-fixed setup with an oblique-plane microscope (OPM).**

**(A) CAD rendering of the rail system carrying the Petri-dish holder and the projection screen.**

**(B) CAD rendering of the tracking and stimulus display setup** (the components below the yellow filter in **Figure 1B**).

**(C) Head-fixed behavioral setup mounted beneath the OPM.**

**(D) Cross-sectional view of the assembled system.**

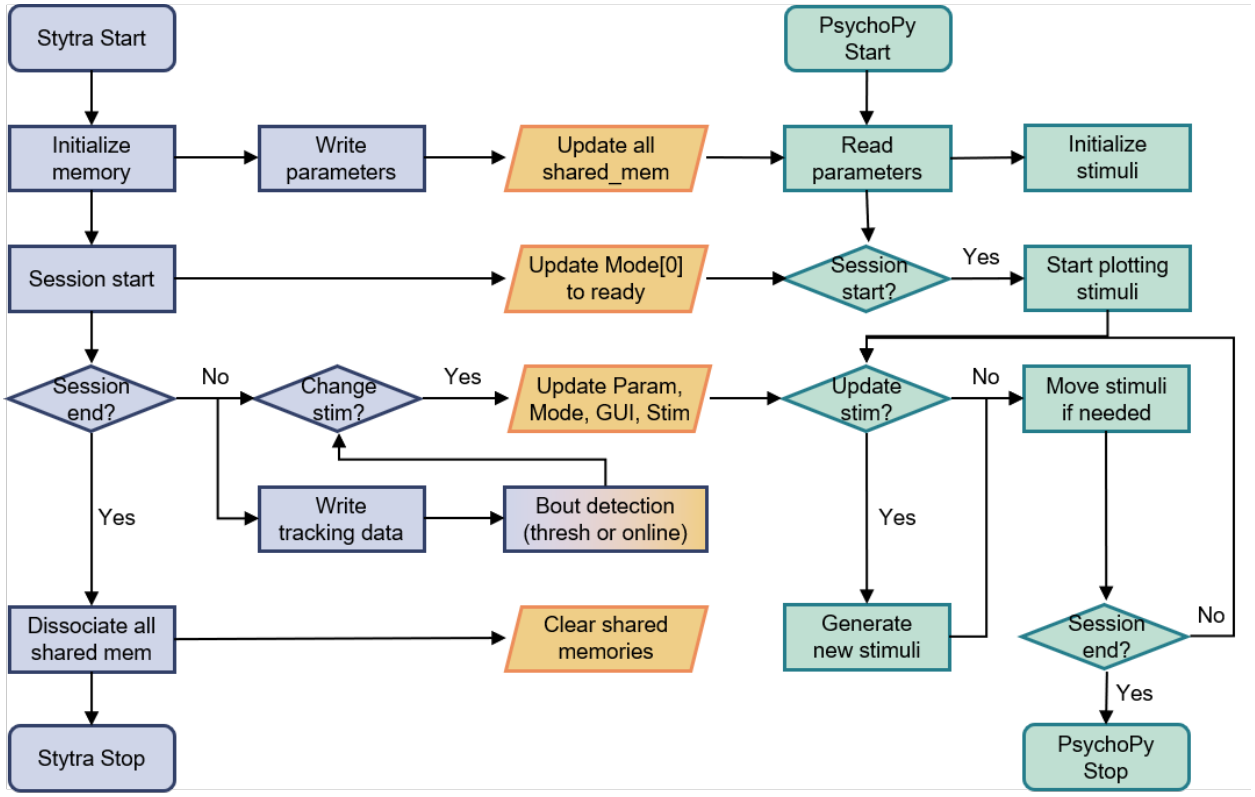


**Figure S2. Inter-process communication (IPC) between Stytra and PsychoPy.**

Schematic of the shared-memory handshake that synchronizes tracking and stimulus generation. Blue blocks denote Stytra processes; yellow blocks denote shared-memory input/output; green blocks denote PsychoPy processes; the blue-to-orange gradient block denotes the real-time bout detector, either the Stytra threshold detector (used to drive closed-loop feedback for all comparative datasets) or the online model (used only for the **Figure S14** real-time cohort). Buffer names and data types are listed in **Table S1**.

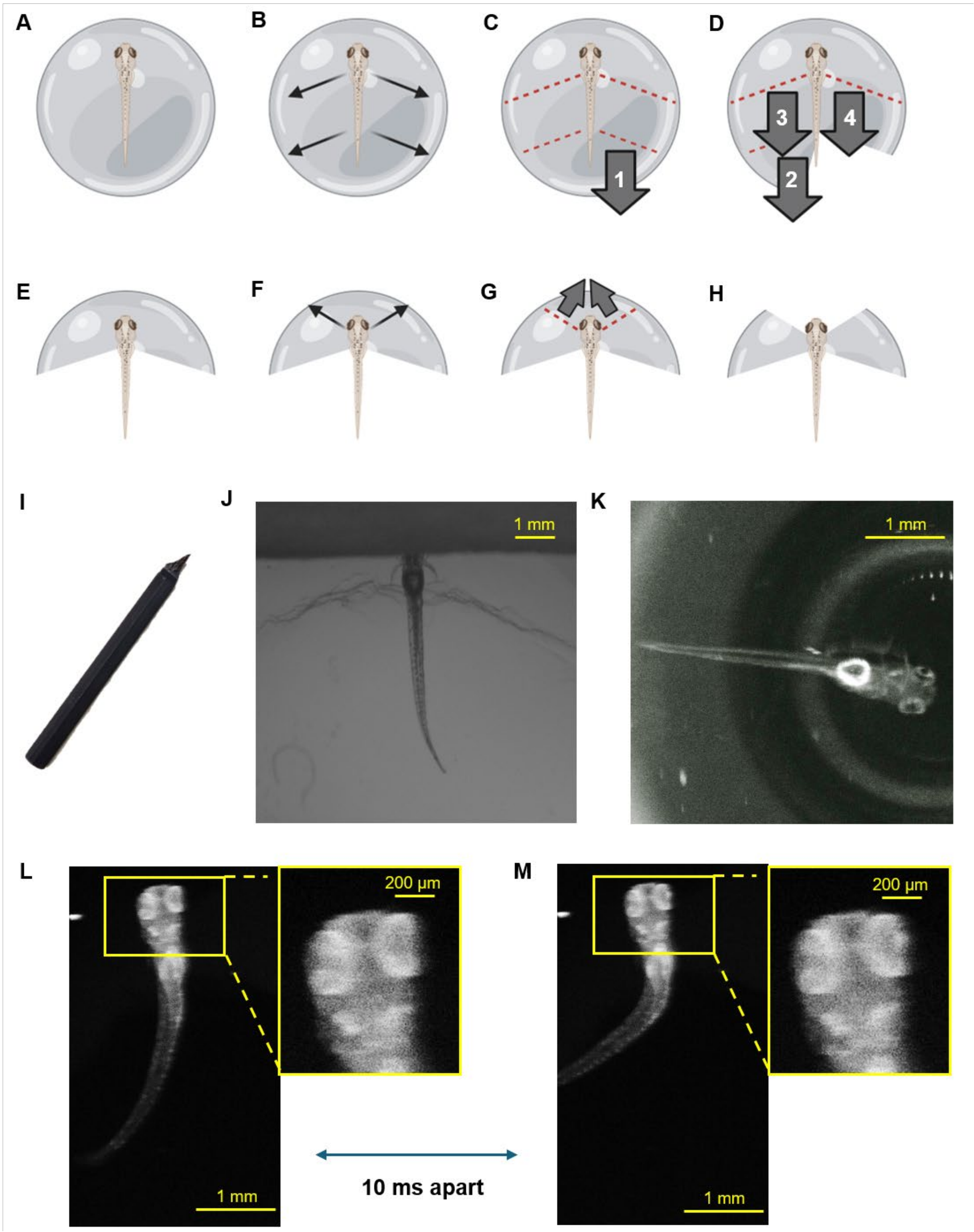


**Figure S3. Agarose mounting protocol.**

**(A-H) Step-by-step mounting procedure used to free the tail and eyes of a head-fixed larva from low-melting-point agarose.**

**(I) The 3D-printed blade tool (0.08 mm tip thickness) used to cut the agarose.**

**(J) A mounted larva viewed through the head-fixed tracking setup, showing the weighing-paper display screen with its aperture.**

**(K) A mounted larva viewed under the OPM.**

**(L-M) Example of eye movement enabled with the mounting protocol.** Eye-position shift between two frames acquired 10 ms apart during the OMR assay, confirming that freeing the eyes preserves eye motility.

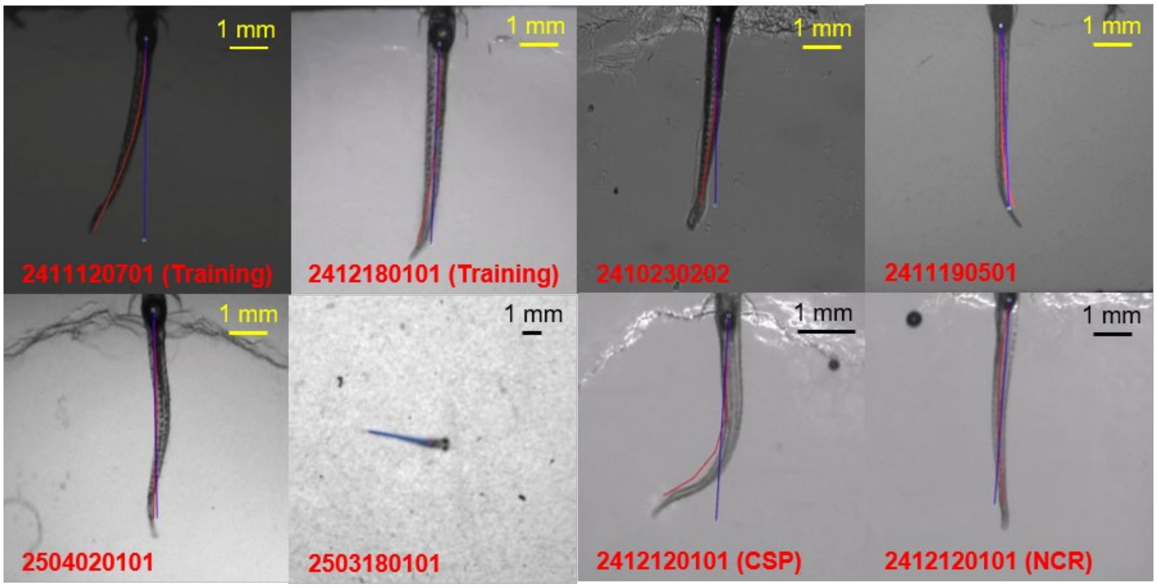


**Figure S4. Tracking quality across imaging conditions and genotypes.**

Representative tracking screenshots for the two training and three testing sessions, spanning the range of lighting conditions in the dataset; the testing set includes lighting conditions absent from the training set. Additional screenshots show free-swimming, casper and nacre sessions, documenting that per-session manual inspection maintained tracking quality across both imaging conditions and pigmentation backgrounds.

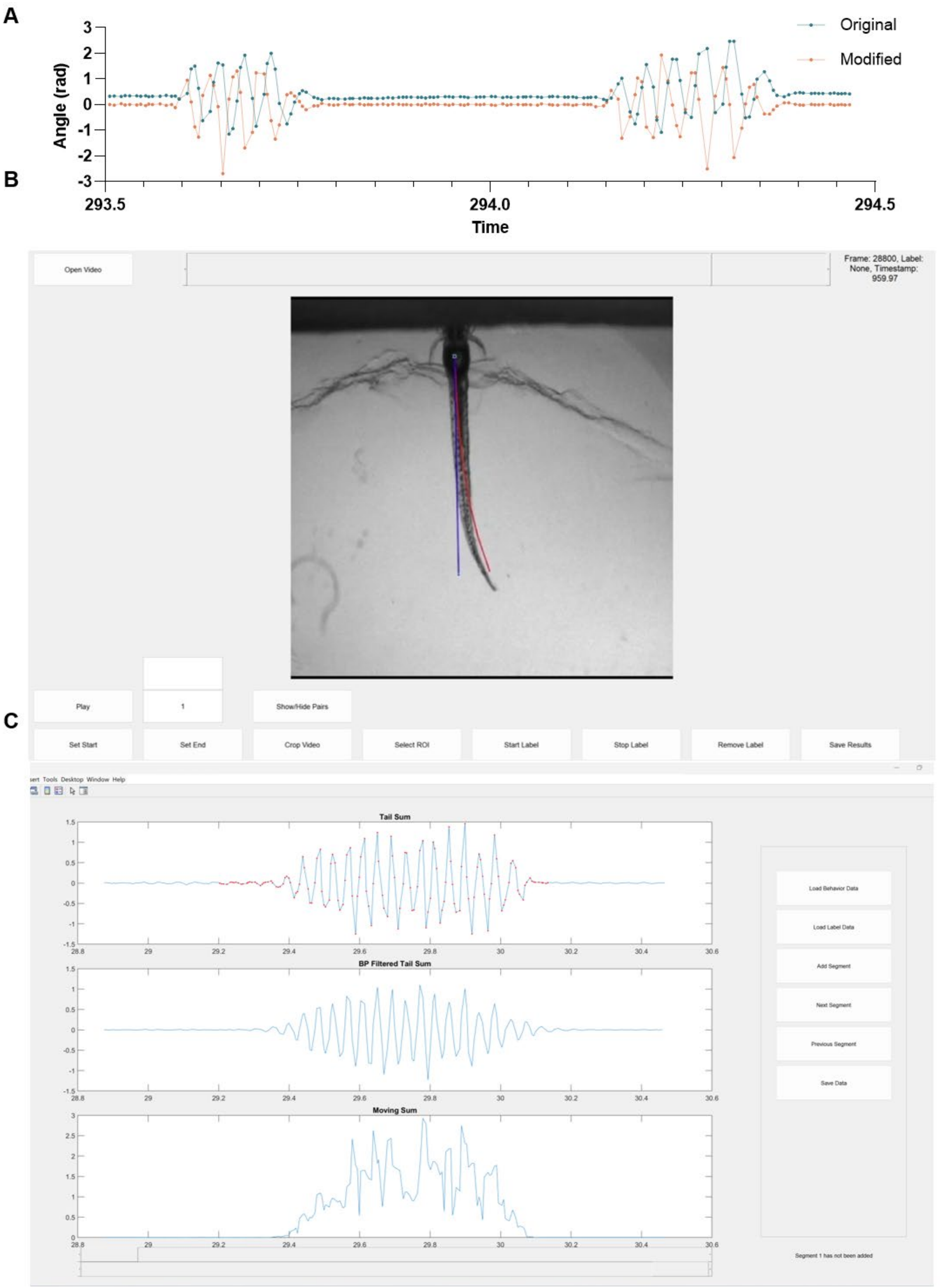


**Figure S5. Two-stage manual annotation interface.**

**(A) Example of the original tail_sum trace from Stytra and the modified version used in this work**. Taking the temporal derivative removes constant offsets and slow drift, effectively acting as a high-pass filter that enhances rapid changes associated with tail motion. Although differentiation introduces a frequency-dependent phase shift, bout kinematics are preserved because both bout onset and offset are detected consistently from the transformed signal.

**(B) Coarse-labeling graphical user interface (MATLAB, *videoAnalysisTool*),** used to review the synchronized Stytra screen recording and save preliminary bout start/end labels; a bout is marked when the distal tail bends laterally relative to the stationary proximal tail. The skeletonized tail segments are shown in red.

**(C) Refined-labeling interface (*boutExtraction*),** used to adjust each bout's onset and offset with a slider while displaying the raw tail_sum trace together with its moving sum of squares and the 20-40 Hz bandpass filtered tail_sum. No amplitude or duration cutoff was applied during labeling.

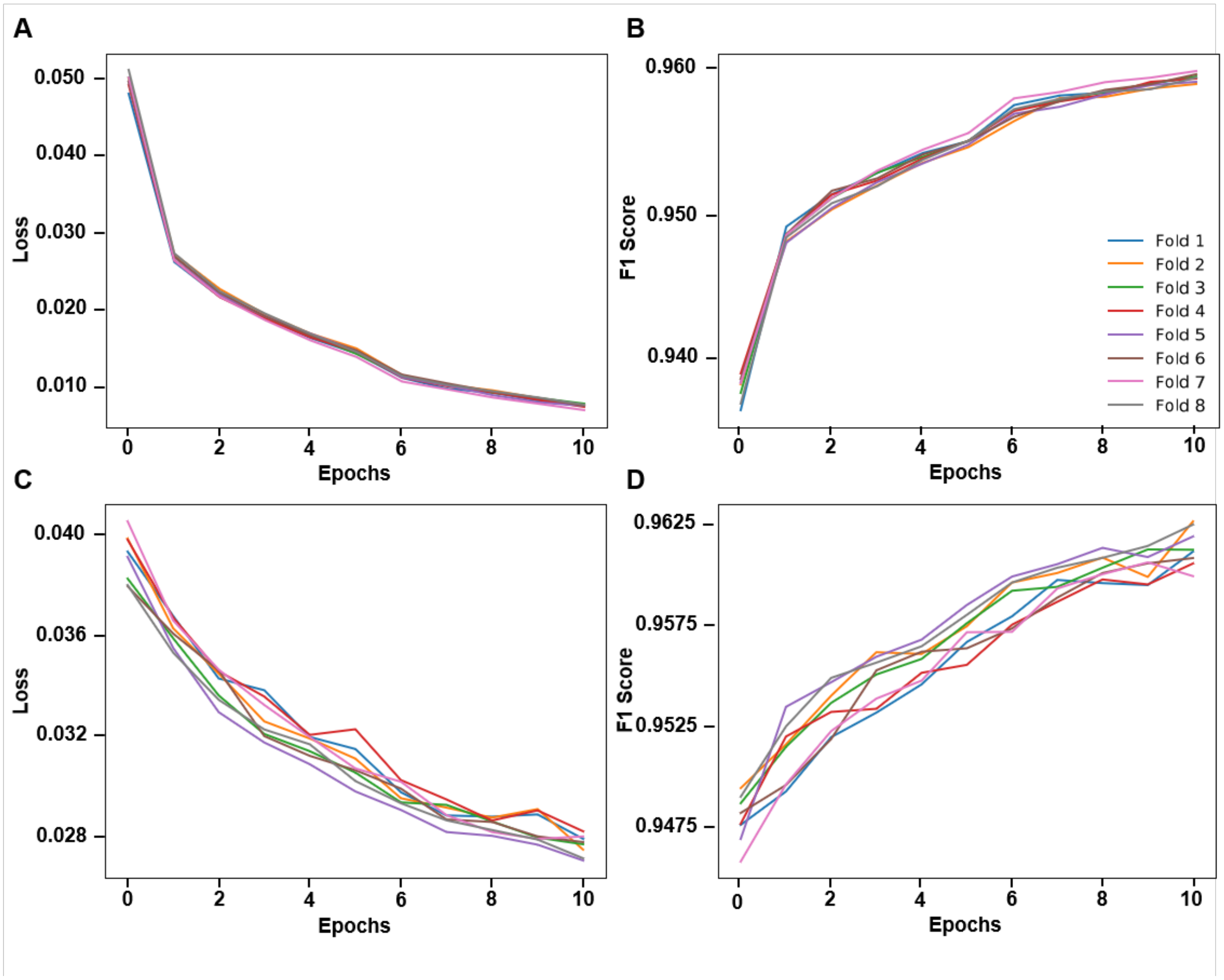


**Figure S6. Training and validation loss curves across 8 folds for offline model. (A)** Offline model training loss. **(B)** Offline model training F1 score. **(C)** Offline model validation loss. **(D)** Offline model validation F1 score.

*Note: color schemes are the same for all four panels.*

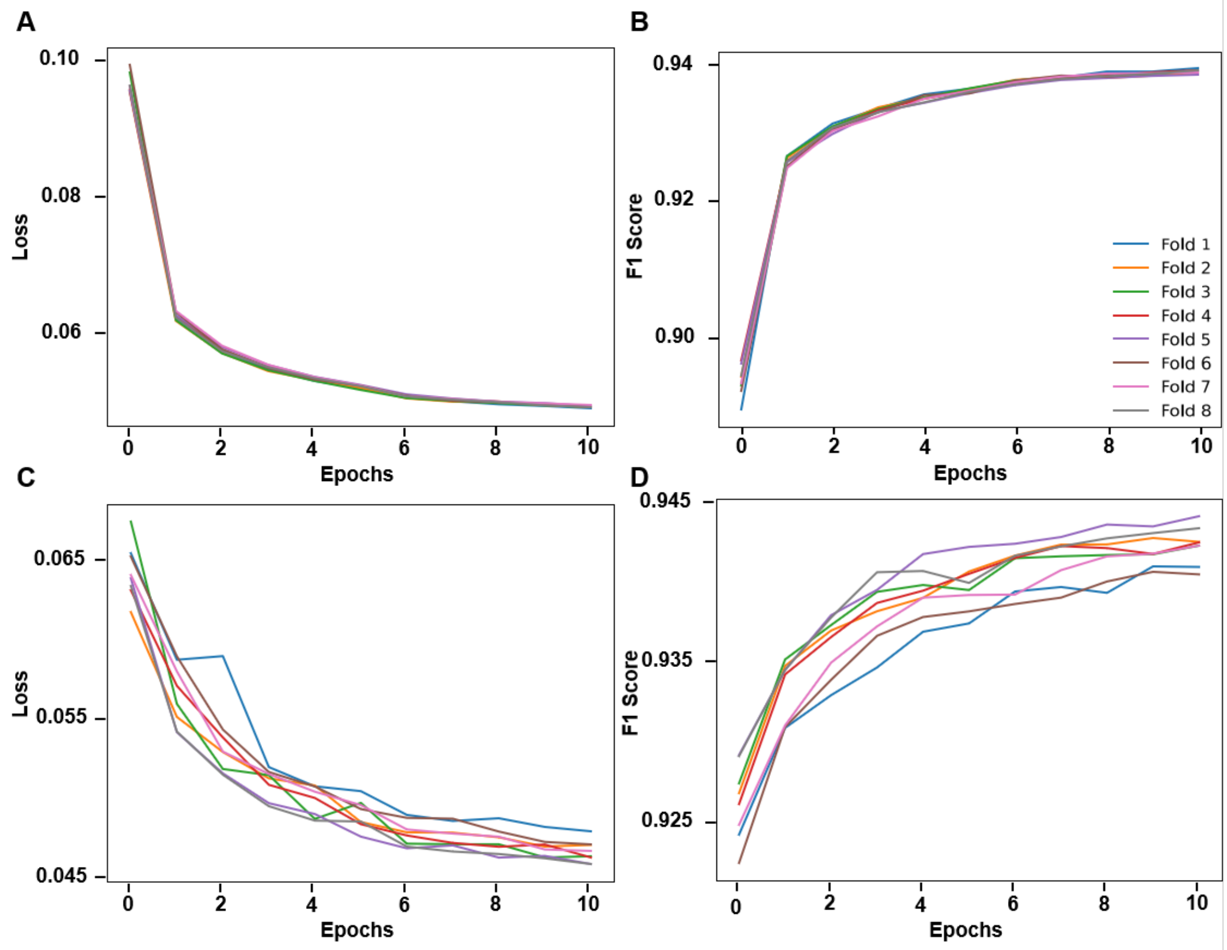


**Figure S7. Training and validation loss curves across 8 folds for online model. (A)** Online model training loss. **(B)** Online model training F1 score. **(C)** Online model validation loss. **(D)** Online model validation F1 score.

*Note: color schemes are the same for all four panels A-D.*

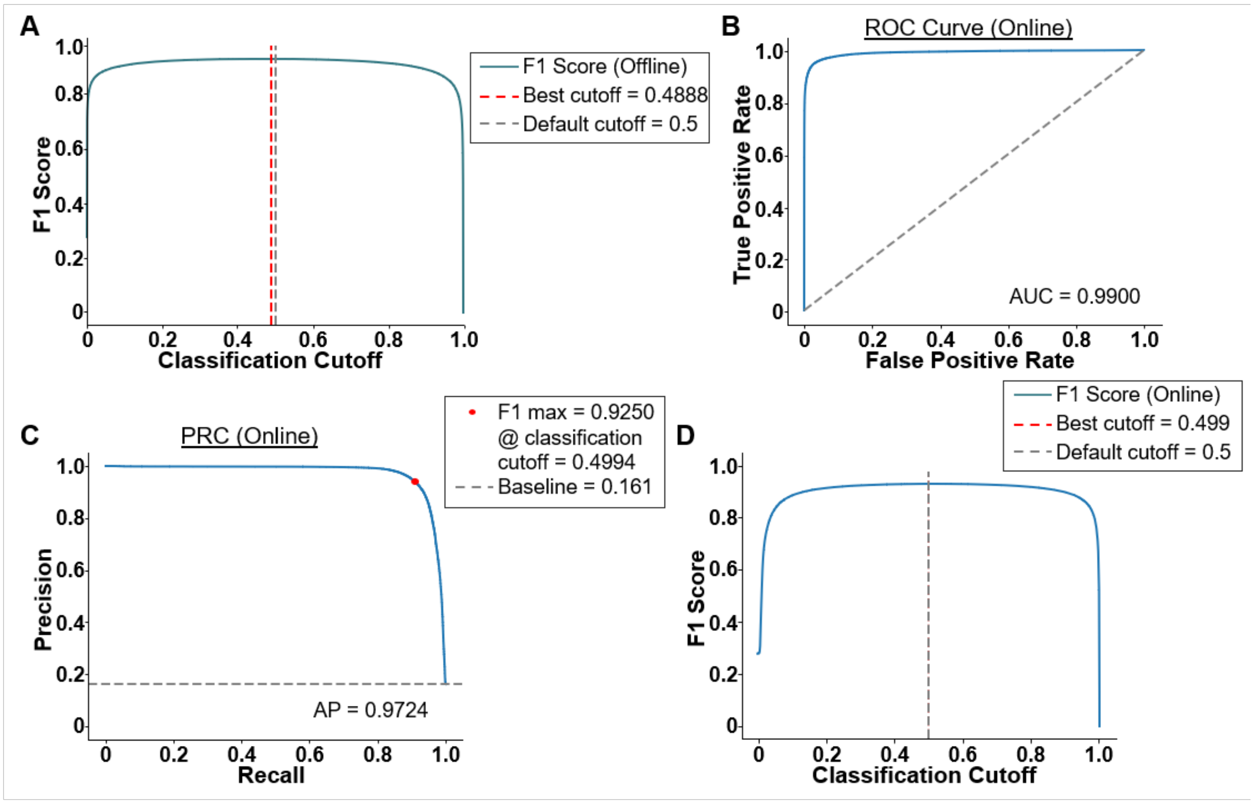


**Figure S8. Classification-cutoff screening and online-model diagnostics.** Head-fixed testing set (N = 6 fish, 6-7 dpf, AB wild-type).

**(A) Offline-model F1 versus classification cutoff**. F1 is maximized at a cutoff of 0.4888, close to the fixed 0.5 cutoff used throughout.

**(B) Online-model receiver-operating-characteristic curve**. AUC = 0.9900.

**(C) Online-model precision-recall curve**. AP = 0.9724 (prevalence baseline = 0.161), with F1 maximized at 0.9250 at a cutoff of 0.4994.

**(D) Online-model F1 versus cutoff**. Best F1 at a cutoff of 0.4994.

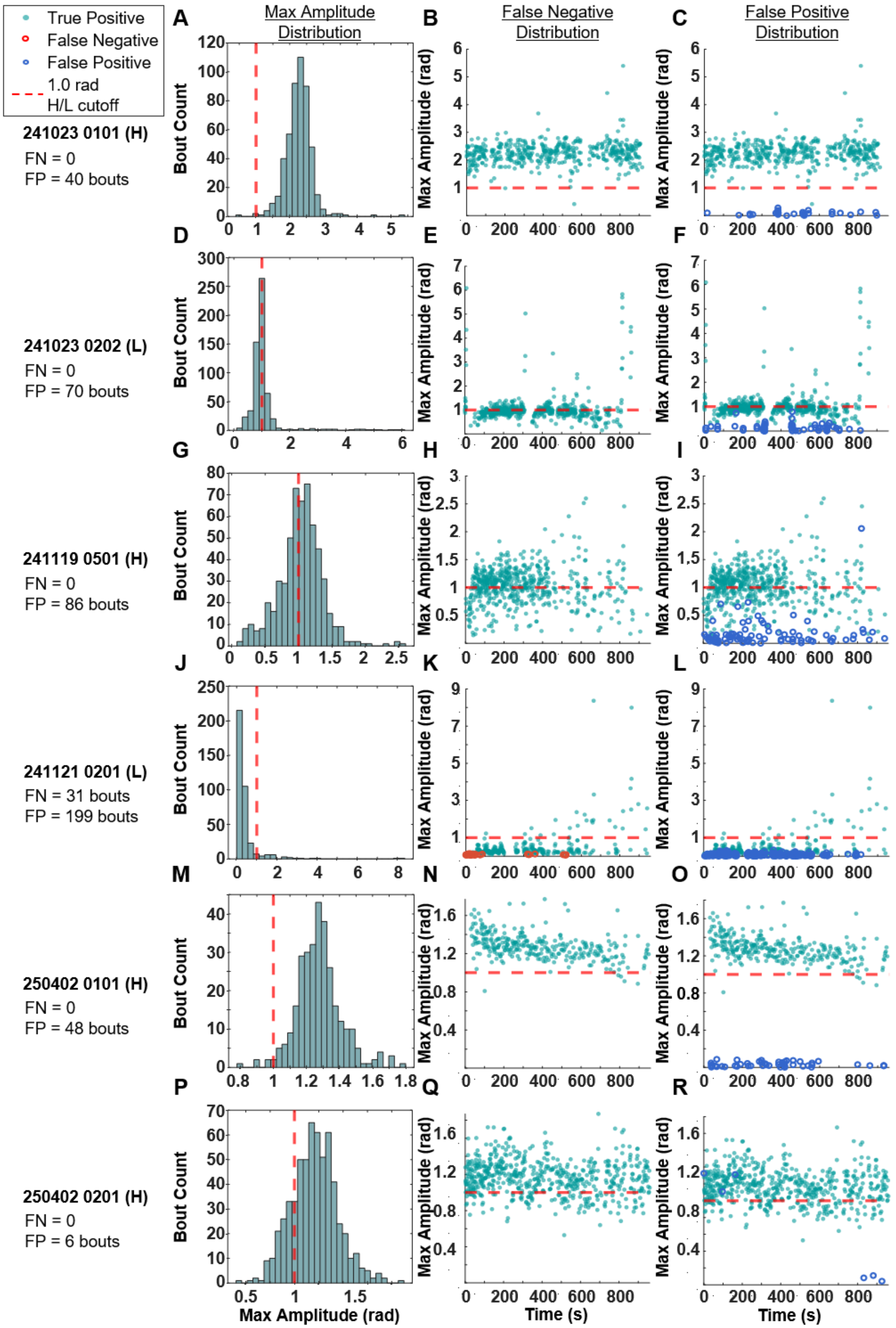
True Positive
False Negative
False Positive
1.0 rad
H/L cutoff
241023 0101 (H)
FN = 0
FP = 40 bouts
241023 0202 (L)
FN = 0
FP = 70 bouts
241119 0501 (H)
FN = 0
FP = 86 bouts
241121 0201 (L)
FN = 31 bouts
FP = 199 bouts
250402 0101 (H)
FN = 0
FP = 48 bouts
250402 0201 (H)
FN = 0
FP = 6 bouts
A
Max Amplitude
Distribution
B
False Negative
Distribution
C
False Positive
Distribution
D
E
F
G
H
I
J
K
L
M
N
O
P
Q
R
Bout Count
Max Amplitude (rad)
Time (s)

**Figure S9. Bout-amplitude distribution and event-level errors over time.**

Head-fixed testing set (N = 6 fish). For each session, a histogram of peak-to-peak bout amplitude is shown together with the temporal distribution of the offline model's false positives (FP) and false negatives (FN), scored at the event (bout) level. A false negative is a manually labeled bout containing no bout label from the offline model; a false positive is a model bout label falling outside any manually labeled bout.

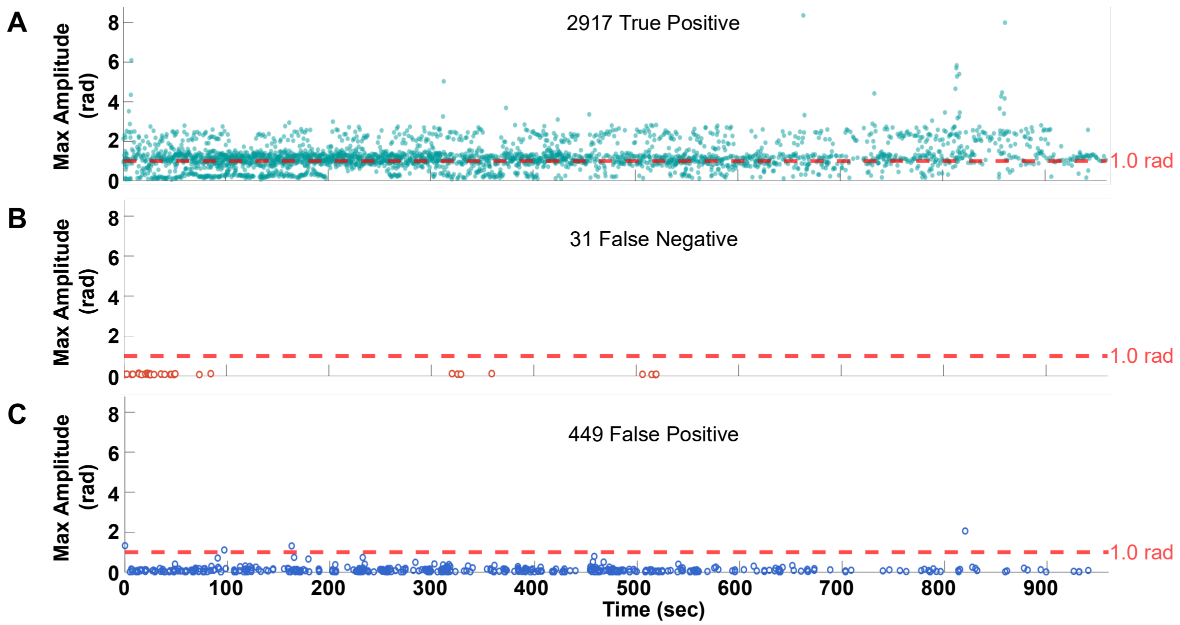


**Figure S10. True-positive, false-negative, and false-positive amplitudes across time.**

**(A) True-positive**, **(B) false-negative**, and **(C) false-positive** bout amplitudes plotted against time within session, aggregated over the all manually labeled fish (N = 14 fish total, 8 from training set and 6 from the testing set) with event-level scoring.

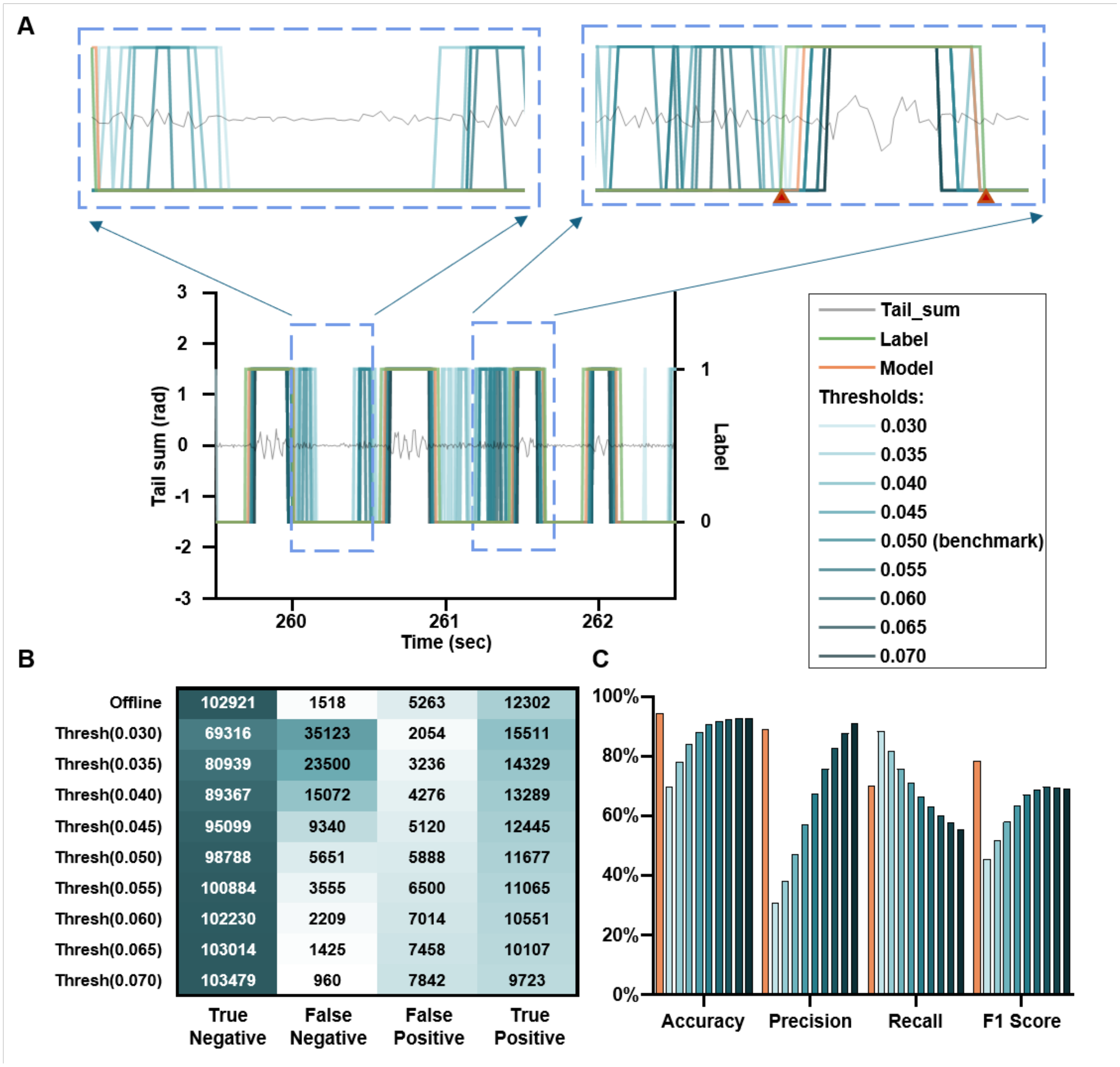


**Figure S11. Threshold-based detection on a low-amplitude session.** Low-amplitude testing session (7.87% of bouts exceed 1.0 rad).

**(A) Performance of the Stytra threshold detector at its default setting (0.050) and at thresholds from 0.030 to 0.070**. Zoomed views show false positives near genuine bouts at low thresholds, whereas at 0.065 and 0.070 the false positives vanish but the onset and offset of manually labeled bouts can no longer be resolved.

**(B-C) Confusion matrices and accuracy, precision, recall, and F1 for the offline model and the threshold detector across the full low-amplitude dataset**. Threshold accuracy plateaus at 0.9244 (0.060), 0.9272 (0.065), and 0.9279 (0.070) versus 0.9444 for the offline model; threshold precision reaches 0.9101 at 0.070 versus 0.8902 offline; threshold recall falls from 0.8831 to 0.5535 as the threshold rises from 0.030 to 0.070, whereas offline recall holds at 0.7004; offline F1 is 0.7839 versus a threshold maximum of 0.6950 (at 0.060). In the traces, gray denotes tail_sum, green manual bouts, orange offline-model bouts, and teal threshold bouts (lighter = lower threshold, darker = higher).

106

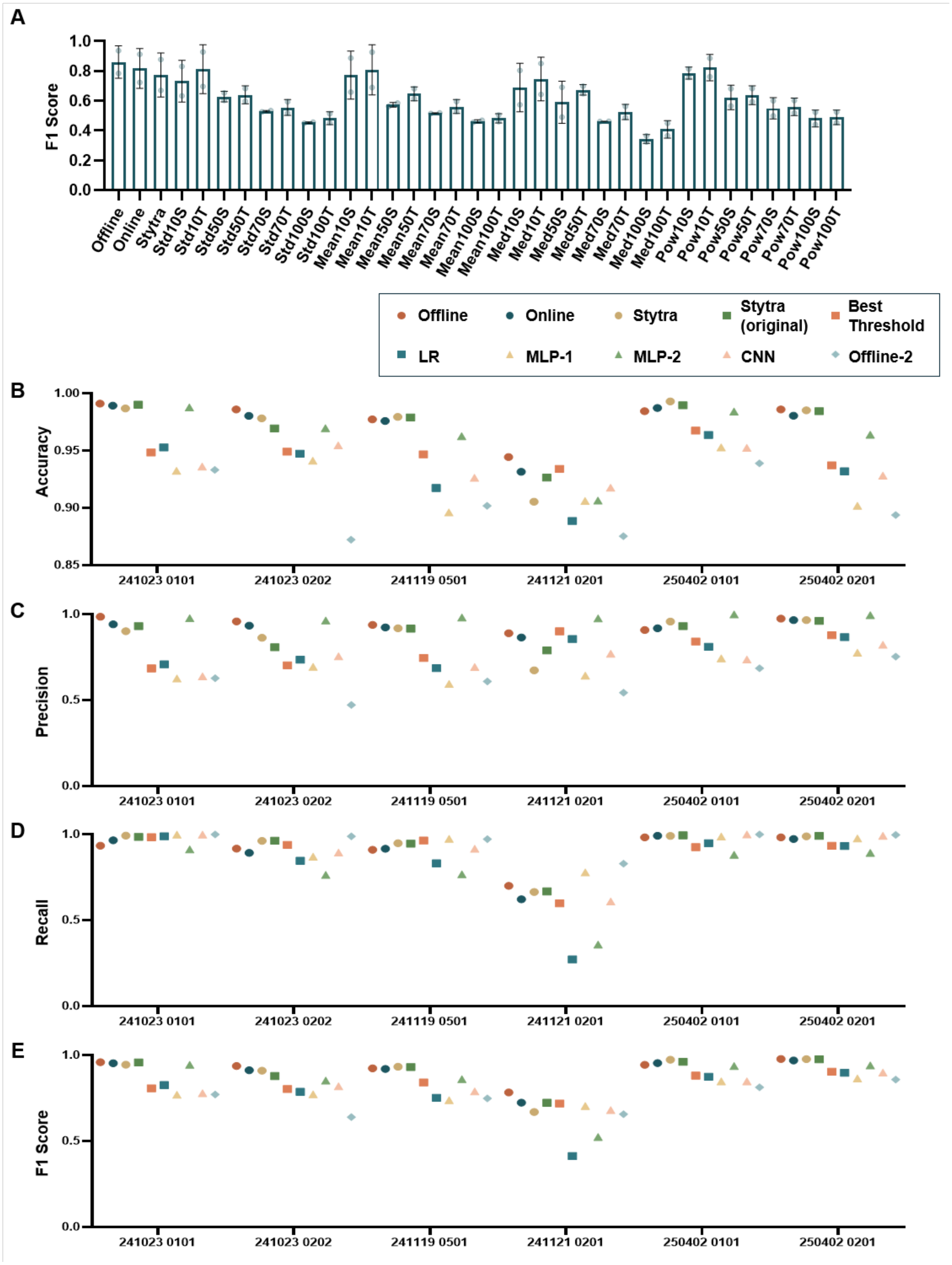


107 **Figure S12. Expanded threshold benchmark and simpler machine-learning baselines.**

**(A) Mean F1 on the two low-amplitude testing sessions for 32 threshold variants**, spanning four statistic families (rolling standard deviation [Std], rolling mean [Mean], rolling median [Med], and moving average of the 20-40 Hz band-pass-filtered tail_sum power [Pow]), four window lengths (10, 50, 70, 100 points), and global static (S) versus per-session-tuned (T) calibration, alongside the offline model, online model, and the native Stytra detector. Static (S) variants were tuned on the training set and applied to the testing set; per-session-tuned (T) variants were calibrated directly on the testing sessions. The per-session-tuned 10-point band-pass-power variant (Pow10T) reached the highest F1 (0.8239) but requires session-level labels; the best static variant (Pow10S; 0.7864) was adopted as the best threshold method for downstream comparisons. The offline (0.8608) and online (0.8183) models exceeded all static variants.

**(B-E) Accuracy, precision, recall, and F1 on the head-fixed testing set (N = 6)** for the offline model, online model, Stytra (modified tail_sum), Stytra (original tail_sum), best threshold method (Pow10S), logistic regression (LR), MLP-1, MLP-2, single-layer CNN, and Offline-2 (offline architecture on six raw tail-segment angles).

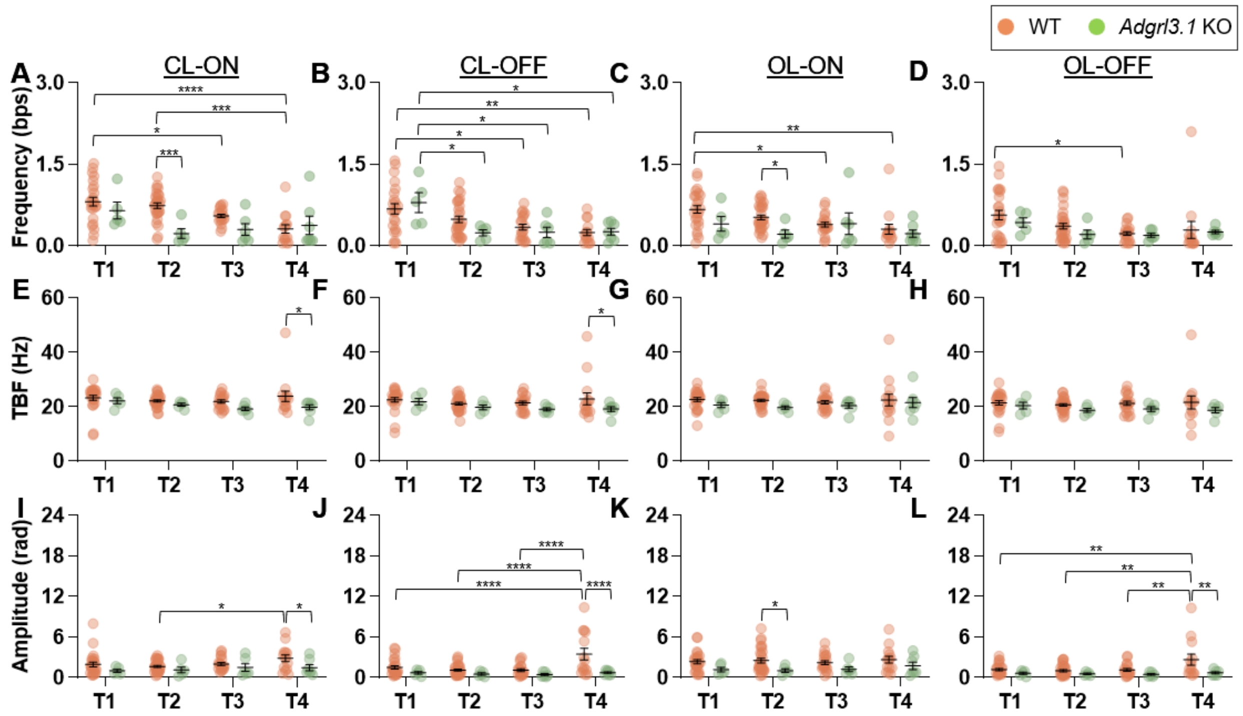


**Figure S13. Additional bout kinematics in the adgrl3.1 knockout.** Head-fixed *adgrl3.1* KO (N = 11, 7 dpf, AB) versus WT AB (N = 55, 6-7 dpf); closed-loop feedback driven by the Stytra threshold detector, with bout labels from the offline model. Two-way ANOVA (strain × trial) with Tukey's post-hoc; ; 95% CIs are of the pairwise mean difference. Panels are arranged by kinematic (rows) and condition CL-ON / CL-OFF / OL-ON / OL-OFF (columns). Bars = mean ± SEM, points = single fish.

**(A-D) Bout frequency.** A strain effect is present at CL-ON ($p = 0.0034$) and OL-ON ($p = 0.0221$) but not CL-OFF ($p = 0.4769$) or OL-OFF ($p = 0.2666$); KO frequency is generally lower than WT. All significant pairwise comparisons: **(A)** at CL-ON, WT > KO at Trial 2 (mean difference 0.5113, 95% CI 0.2242-0.7983, $p = 0.0006$), with WT declining across trials (WT T1 > T3: mean difference 0.2601, 95% CI 0.02095-0.4992, $p = 0.0274$; T1 > T4: mean difference 0.4987, 95% CI 0.2369-0.7606, $p < 0.0001$; T2 > T4: mean difference 0.4265, 95% CI 0.1716-0.6814, $p = 0.0002$); **(B)** at CL-OFF, both strains declined across trials (WT T1 > T3: mean difference 0.3355, 95% CI 0.08329-0.5878, $p = 0.0042$; WT T1 > T4: mean difference 0.4359, 95% CI 0.1601-0.7117, $p = 0.0004$; KO T1 > T2: mean difference 0.5567, 95% CI 0.04204-1.071, $p = 0.0286$; KO T1 > T3: mean difference 0.5467, 95% CI 0.05395-1.039, $p = 0.0235$; KO T1 > T4: mean difference 0.5348, 95% CI 0.05831-1.011, $p = 0.0214$); **(C)** at OL-ON, WT > KO at Trial 2 (mean difference 0.3061, 95% CI 0.03130-0.5808, $p = 0.0294$), with WT declining across trials (WT T1 > T3: mean difference 0.2788, 95% CI 0.04995-0.5077, $p = 0.0103$; T1 > T4: mean difference 0.3616, 95% CI 0.1109-0.6122, $p = 0.0016$); **(D)** and at OL-OFF, WT T1 > T3 (mean difference 0.3410, 95% CI 0.07520-0.6067, $p = 0.0062$).

**(E-H) Tail-beat frequency**. A strain effect is present at CL-ON ($p = 0.0132$), CL-OFF ($p = 0.0373$), and OL-OFF ($p = 0.0487$) but not OL-ON ($p = 0.0731$). Significant pairwise: **(E)** WT > KO at Trial 4 under CL-ON (mean difference 4.077, 95% CI 0.5336-7.621, $p = 0.0246$) and **(F)** CL-OFF (mean difference 3.751, 95% CI 0.04381-7.458, $p = 0.0474$); no pairwise comparison reached significance at OL-ON **(G)** or OL-OFF **(H)**.

**(I-L) Maximum amplitude.** A strain effect is present in all four conditions (CL-ON $p = 0.0082$; CL-OFF $p = 0.0005$; OL-ON $p = 0.0026$; OL-OFF $p = 0.0030$), with KO amplitudes generally lower than WT. All significant pairwise comparisons: **(I)** at CL-ON, WT > KO at Trial 4 (mean difference 1.370, 95% CI 0.1757-2.564, $p = 0.0250$), with WT amplitude rising from Trial 2 to Trial 4 (WT T2 < T4: mean difference -1.189, 95% CI -2.301 to -0.07719, $p = 0.0312$);

**(J)** at CL-OFF, WT > KO at Trial 4 (mean difference 2.743, 95% CI 1.442-4.044, $p < 0.0001$) with a WT-specific Trial 4 amplitude rise (WT T1 < T4: mean difference -1.959, 95% CI -3.214 to -0.7040, $p = 0.0005$; T2 < T4: mean difference -2.361, 95% CI -3.573 to -1.150, $p < 0.0001$; T3 < T4: mean difference -2.359, 95% CI -3.663 to -1.055, $p < 0.0001$); **(K)** at OL-ON, WT > KO at Trial 2 (mean difference 1.470, 95% CI 0.005810-2.935, $p = 0.0491$); **(L)** and at OL-OFF, WT > KO at Trial 4 (mean difference 1.928, 95% CI 0.7057-3.149, $p = 0.0023$) with a WT Trial 4 amplitude rise (WT T1 < T4: mean difference -1.430, 95% CI -2.553 to -0.3073, $p = 0.0066$; T2 < T4: mean difference -1.605, 95% CI -2.699 to -0.5109, $p = 0.0013$; T3 < T4: mean difference -1.507, 95% CI -2.681 to -0.3339, $p = 0.0061$).

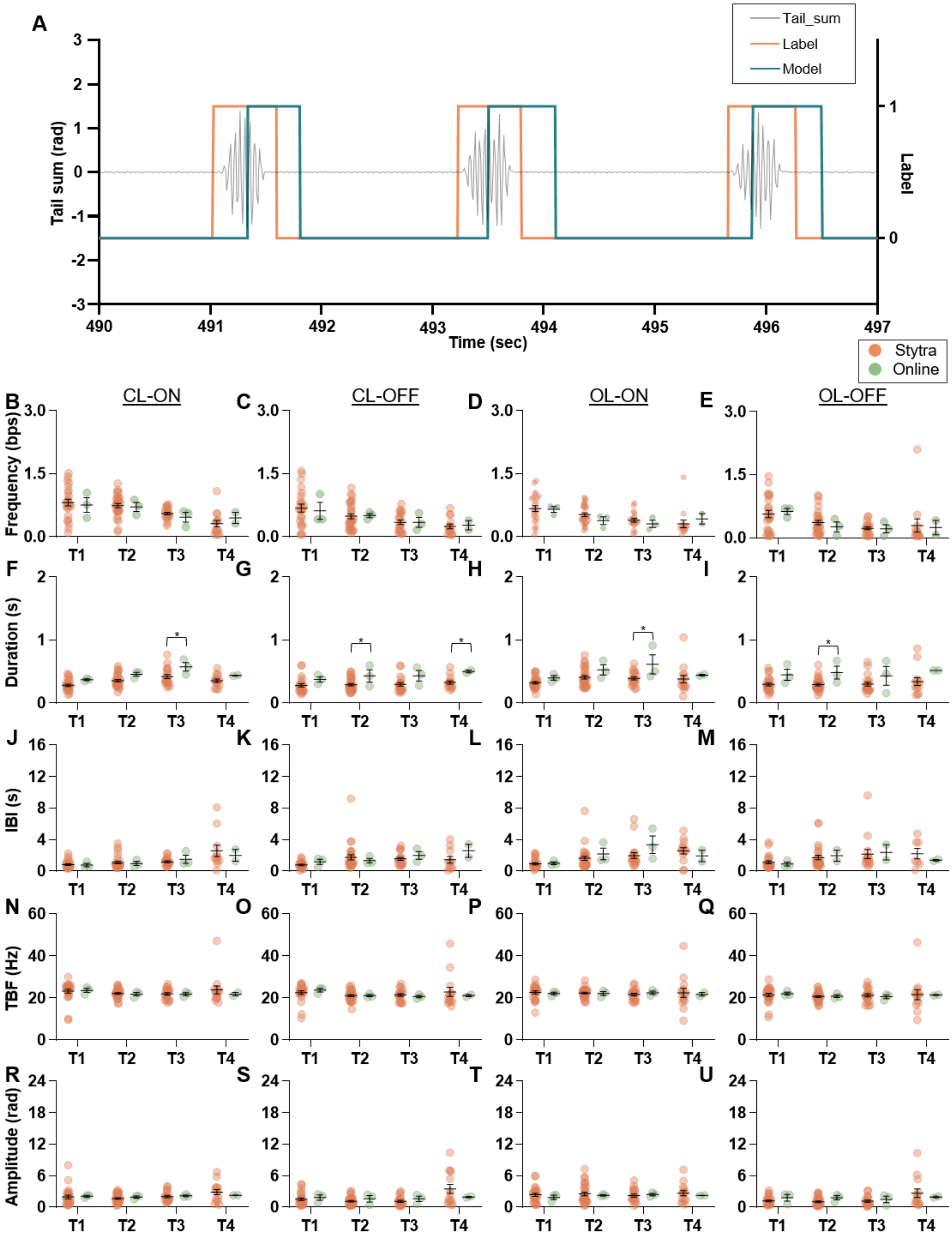


**Figure S14. Real-time application of the online model and comparison of closed-loop drivers.**

**(A) Example real-time tail_sum trace and online-model detection envelope from one 5-dpf WT AB larva**. Mean bout-onset latency was 209.3, 211.3, and 218.8 ms across three labeled sessions.

**(B-U) Bout kinematics of the online-driven cohort (N = 5, 5 dpf; closed-loop feedback driven by the online model) versus the Stytra-driven cohort (N = 55)** under two-way ANOVA (driver cohort × trial): frequency **(B-E)**, duration **(F-I)**, interbout interval **(J-M)**, tail-beat frequency **(N-Q)**, and maximum amplitude **(R-U)**. Trial effects of the Stytra driven are reported in **Figure 5** and thus omitted here. Only bout duration differed between drivers, lengthening in the online-driven cohort across all four conditions (cohort main effect: CL-ON p = 0.0023; CL-OFF p = 0.0004; OL-ON p = 0.0120; OL-OFF p = 0.0006); the difference reached significance at individual trials, **(F)** CL-ON Trial 3 (Stytra < online: mean difference -0.1542 s, 95% CI -0.2829 to -0.02546, p = 0.0195); **(G)** CL-OFF Trial 2 (Stytra < online: mean difference -0.1398 s, 95% CI -0.2777 to -0.001932, p =0.0469) and Trial 4 (Stytra < online: mean difference -0.1749 s, 95% CI -0.3465 to -0.003391, p = 0.0457); **(H)** OL-ON Trial 3 (Stytra < online: mean difference -0.2233 s, 95% CI -0.4001 to -0.04644, p = 0.0139); and **(I)** OL-OFF Trial 2 (Stytra < online: mean difference -0.1899 s, 95% CI -0.3587 to -0.02114, p = 0.0279). Frequency, interbout interval, tail-beat frequency, and maximum amplitude were unaffected (all p > 0.3).

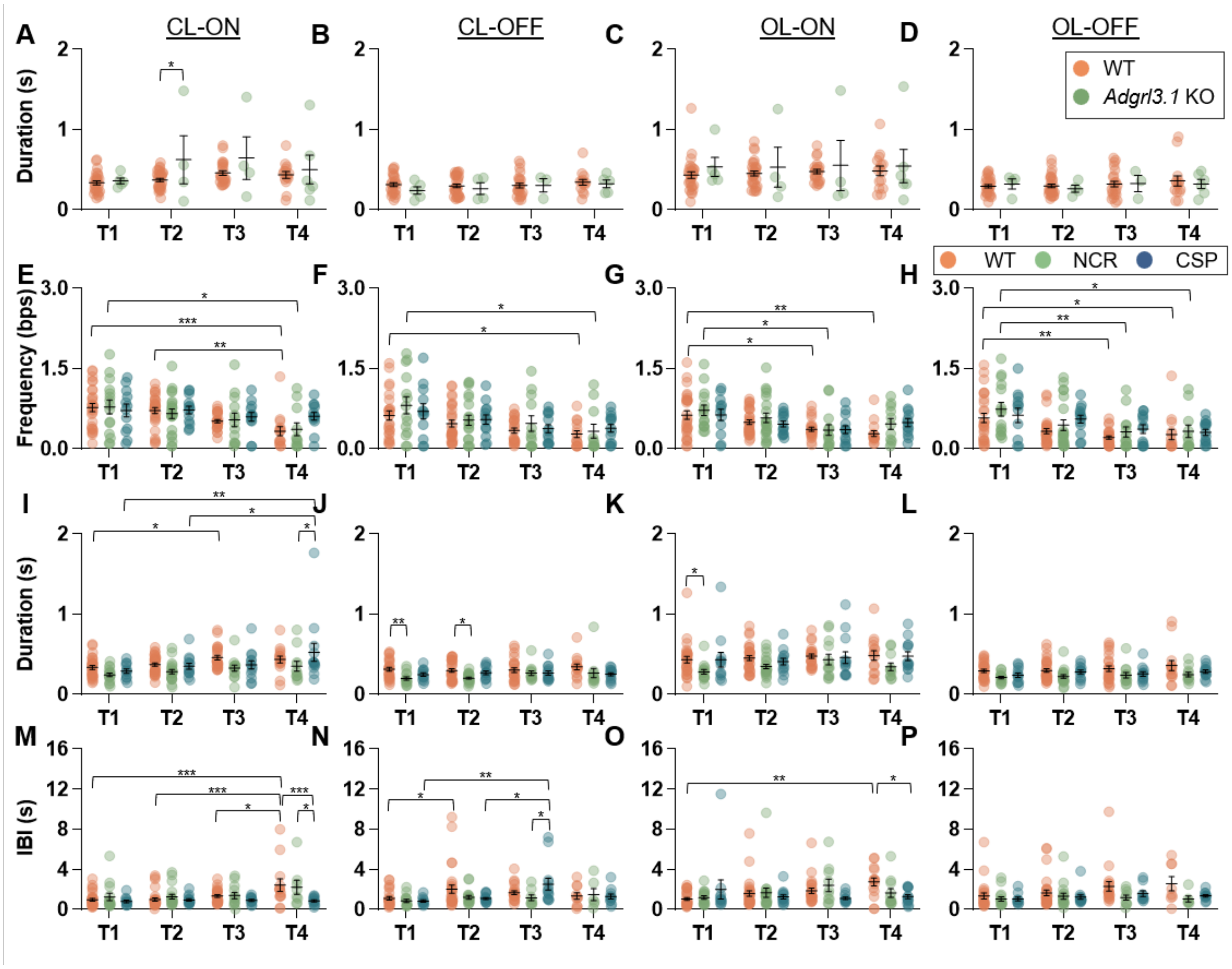


**Figure S15. Bout kinematics under the best threshold method as a companion to the offline-model results.** Same head-fixed cohorts and statistical design as **Figures 4** and **5**, re-analyzed with the best threshold method (Pow10S) in place of the offline model; two-way ANOVA with Tukey's post-hoc. **(A-D) Bout duration of *adgrl3.1* KO versus WT** (companion to **Figure 4 A-D**). **(E-H) Bout frequency of WT, nacre, and casper** (companion to **Figure 5 A-D**). **(I-L) Bout duration of WT, nacre, and casper** (companion to **Figure 5 E-H**). **(M-P) Interbout interval of WT, nacre, and casper** (companion to **Figure 5 I-L**). Across all comparisons the two detectors agree on the large majority of contrasts and differ on 22 (offline-only on 13, threshold-only on 9): for *adgrl* duration the offline model reaches significance on 2 contrasts not reached by the threshold method (**Table S9**); for frequency the offline model reaches 5 and the threshold method 1 (**Table S10**); for duration the two diverge on 4 contrasts each (**Table S11**); for interbout interval the offline model reaches 2 and the threshold method 4 (**Table S12**).

**Table S1. Shared-memory variables for Stytra-PsychoPy inter-process communication.** Buffer names, data types, and command codes for the shared-memory channel that relays tracking data and stimulus commands between Stytra and PsychoPy (**Figure S1**).

| **Buffer Name [index, bounds are inclusive]** | **Data stored (data type)** | **Possible codons/values and explanation** |
|---|---|---|
| **Mode[0]** | Current state of experiment (int) | 0: Stytra has not yet started<br>1: open loop started<br>2: open loop standby<br>3: closed loop started<br>4: closed loop standby |
| **Stim[0]** | Type of stimulus (int) | 0: sinusoidal<br>1: square<br>2: sawtooth<br>3: triangular<br>4: continuous leopard background |
| **Stim[1]** | Current state of moving grating (int) | 0: during grating pause<br>1: during moving phase<br>2: reset phase |
| **GUI[0:3]** | self.t_pre from Stytra (float32) | Duration of grating pause in seconds |
| **GUI[4:7]** | self.t_move from Stytra (float32) | Duration of grating movement in seconds |
| **GUI[8:11]** | Grating cycle (float32) | Number of cycles within the display window |
| **GUI[12:15]** | Grating base speed (float32) | The base speed of grating in pixels per second in open loop (or the speed of grating movement when the fish is not moving in closed-loop) |
| **Update[0]** | Current state of the fish in closed-loop experiments (int) | 0: no change<br>1: Fish has moved and the speed and/or the angle of the grating need to be updated |
| **Param[0:3]** | N/A | |
| **Param[4:7]** | self.vel from Stytra (float32) | Speed of grating with closed-loop feedback from intended velocity of the fish, in pixels per second |
| **Param[8:11]** | Theta from Stytra (float32) | The angle of grating in degrees |
| **Param[12:15]** | Duration from Stytra (float32) | The duration of each session in seconds |
| **SEQ[0:119]** | Tail_sum data (float32) | Tail_sum in radians from the previous 30 tracking results for real-time bout detection |

**Table S2. Composition and annotation reliability of the training and testing sets.** Per-session fish identifier, amplitude type (H = high, L = low), percentage of bouts exceeding 1.0 rad maximum peak-to-peak amplitude, and annotation agreement. Intra-rater agreement (Cohen's κ and percent agreement) is given for all training sessions and 4 of the testing sessions; inter-rater agreement (κ and percent agreement, one entry per additional annotator, N = 3 additional annotators) is given for the re-labeled testing sessions. **(A) Training set**: N = 8 fish (3743 bouts); **(B) testing set**: N = 6 fish (2948 bouts).

| Fish ID (Type) | Percent High | Intra-rater κ/ percent agreement |
|---|---|---|
| **241023 0102 (H)** | 100% | 0.8235/ 0.9371 |
| **241023 0201 (H)** | 83.40% | 0.7315/ 0.8973 |
| **241105 0401 (H)** | 92.44% | 0.7429/ 0.9140 |
| **241112 0701 (H)** | 98.12% | 0.7977/ 0.9562 |
| **241118 0201 (L)** | 27.40% | 0.6704/ 0.9826 |
| **241119 0101 (L)** | 4.83% | 0.7334/ 0.9206 |
| **241203 0601 (L)** | 6.98% | 0.7266/ 0.9333 |
| **241218 0101 (L)** | 0.00% | 0.7119/ 0.9434 |

| Fish ID (Type) | Percent High | Intra-rater κ/ percent agreement | Inter-rater(s) κ/percent agreement | |
|---|---|---|---|---|
| **241023 0101 (H)** | 99.80% | 0.7428/ 0.9365 | 0.8647/ 0.9698 | 0.6546/ 0.9064 |
| **241023 0202 (L)** | 20.97% | 0.7240/ 0.9378 | 0.8329/ 0.9614 | N/A |
| **241119 0501 (H)** | 56.11% | 0.7382/ 0.9209 | N/A | N/A |
| **241121 0201 (L)** | 7.87% | N/A | 0.7067/ 0.9252 | 0.6132/ 0.9315 |
| **250402 0101 (H)** | 98.37% | 0.7746/ 0.9378 | 0.9094/ 0.9779 | N/A |
| **250402 0201 (H)** | 80.79% | N/A | 0.9459/ 0.9783 | N/A |

**Table S3. Features and their extraction methods for the offline and online models.** The offline model contains an additional BiLSTM layer, serving as a *post hoc* bout detection method when computational time is not a constraint. To enhance prediction speed and to enable real-time bout prediction, a modified online model includes the six features with highest permutation importance, and the feature extraction methods are adjusted to accelerate the computation. Numpy functions used for the offline model and the mathematical expression used for the online model are provided for reference. For the offline model, max and min values are extracted for the rolling window of the absolute values of the raw traces, while in the online model it is the max and min values of the raw trace.

| Feature | Abbrev. | Definition in offline model *(post hoc)* | Definition in online model (real-time) |
|---|---|---|---|
| **Raw data** | *tail_sum* | The modified (temporal derivative) tail_sum data from tracking | $f_1(i) = \theta_i$<br>• The modified tail_sum data at index i of the segment |
| **Sum of squared values** | *sq* | Calculated as the sum of np.square values within a rolling window of size feature_window | $f_2(i) = \sum_{j=0}^{m-1} \theta_{i+j}^2$<br>• m: feature_window |
| **Mean of absolute values** | *mean* | Calculated as the mean of np.abs values within a rolling window of size feature_window | $f_3(i) = \frac{1}{m}\sum_{j=0}^{m-1} \lvert\theta_{i+j}\rvert$<br>• m: feature_window |
| **First derivative of the raw data** | *speed* | Calculated with np.gradient on the raw data within the rolling window | |
| **Maximum** | *max* | Calculated with np.max on the np.abs values within the rolling window | $f_4(i) = \max\{\theta_{i+j}\},\ j \in [0,\ m]$<br>• m: feature_window |
| **Minimum** | *min* | Calculated with np.min on the np.abs values within the rolling window | $f_5(i) = \min\{\theta_{i+j}\},\ j \in [0,\ m]$<br>• m: feature_window |
| **Mean** | *roll_ mean* | Calculated with np.mean on the raw data within the rolling window | $f_6(i) = \frac{1}{m}\sum_{j=0}^{m-1} \theta_{i+j}$<br>• m: feature_window |
| **Gradient of sum of squared values** | *sq_grad* | Calculated with np.gradient on the np.square values | |
| **Sum of squared values of filtered data** | *filt_sq* | Calculated as the sum of np.square values for $4^{th}$ order Butterworth BPF (20-40 Hz) filtered raw data | |

Table S4. Permutation feature importance (PFI) scores for the presented versions of the offline model and the online model. Definitions and derivation of the features are listed in Table S3.

| **Offline Feature** | **Score** | **Online Feature** | **Score** |
|---|---|---|---|
| *tail_sum* | 0.111 | *tail_sum* | 0.346 |
| *sq* | 0.066 | *sq* | 0.217 |
| *mean* | 0.398 | *mean* | 0.438 |
| *speed* | 0.008 | [redacted] | |
| *max* | 0.018 | *max* | 0.218 |
| *min* | 0.051 | *min* | 0.367 |
| *roll_ mean* | 0.196 | *roll_ mean* | 0.411 |
| *sq_grad* | 0.026 | [redacted] | |
| *filt_sq* | 0.101 | | |

Table S5. Permutation feature-importance scores for the initial feature pool. PFI for the full pre-curation feature pool (raw and band-pass-filtered variants); features retained in the final offline model are highlighted.

| Feature | Permutation Importance | Feature | Permutation Importance |
|---|---|---|---|
| **Tail_sum (raw data)** | 0.1180 | **Tail_sum (after BPF*)** | 0.0022 |
| **Variance** | 0.0053 | **BPF Variance** | 0.0012 |
| **Sum of SQ** | 0.0940 | **BPF Sum of SQ** | 0.2151 |
| **Mean of Abs Value** | 0.4011 | **BPF MAV** | 0.0010 |
| **First derivative (speed)** | 0.0094 | **BPF 1st derivative** | 0.0020 |
| **Max** | 0.0175 | **BPF max** | 0.0043 |
| **Rolling mean** | 0.1872 | **BPF rolling mean** | 0.0007 |
| **Derivative of Sum SQ** | 0.0180 | **BPF derivative Sum SQ** | 0.0006 |
| **Min** | 0.0116 | **BPF min** | 0.0003 |

Table S6. Accuracy, precision, recall (sensitivity), and F1 score for the Stytra threshold-based method, offline model, and online model evaluated on the testing sets. Dataset labels (H for high amplitude, L for low amplitude) indicate whether more than 50% of bouts exceed a maximum tail( sum.amplitude of 1.0 rad, followed by percent of high amplitude bouts. For each metric, the rows correspond to the Stytra threshold-based method, the offline model, and the online model, respectively. Mean values and their standard errors (sem) are reported for all testing datasets, as well as separately for the high- and low-amplitude subsets. Average scores that improve by more than 1% relative to the threshold-based benchmark are highlighted in red, whereas decreases greater than 1% are highlighted in green.

| Fish ID (Type, percent high) | Method | Accuracy | Precision | Recall | F1 Score |
|---|---|---|---|---|---|
| **241023 0101** (H, 99.80%) | Stytra | 0.9869 | 0.9014 | 0.9931 | 0.9450 |
| | Offline | 0.9911 | 0.9863 | 0.9338 | 0.9594 |
| | Online | 0.9894 | 0.9414 | 0.9659 | 0.9535 |
| **241023 0202** (L, 20.97%) | Stytra | 0.9783 | 0.8637 | 0.9623 | 0.9103 |
| | Offline | 0.9860 | 0.9586 | 0.9174 | 0.9376 |
| | Online | 0.9804 | 0.9340 | 0.8923 | 0.9127 |
| **241119 0501** (H, 56.11%) | Stytra | 0.9795 | 0.9186 | 0.9483 | 0.9332 |
| | Offline | 0.9773 | 0.9380 | 0.9100 | 0.9238 |
| | Online | 0.9760 | 0.9238 | 0.9163 | 0.9201 |
| **241121 0201** (L, 7.87%) | Stytra | 0.9054 | 0.6739 | 0.6648 | 0.6693 |
| | Offline | 0.9444 | 0.8902 | 0.7004 | 0.7839 |
| | Online | 0.9316 | 0.8650 | 0.6224 | 0.7239 |
| **250402 0101** (H, 98.37%) | Stytra | 0.9930 | 0.9580 | 0.9908 | 0.9741 |
| | Offline | 0.9845 | 0.9082 | 0.9826 | 0.9439 |
| | Online | 0.9873 | 0.9190 | 0.9922 | 0.9542 |
| **250402 0201** (H, 80.79%) | Stytra | 0.9853 | 0.9665 | 0.9889 | 0.9775 |
| | Offline | 0.9860 | 0.9745 | 0.9822 | 0.9784 |
| | Online | 0.9806 | 0.9666 | 0.9736 | 0.9701 |
| **mean** | Stytra | 0.9705 | 0.8761 | 0.9234 | 0.8984 |
| | Offline | 0.9782 | 0.9426 | 0.9044 | 0.9212 |
| | Online | 0.9742 | 0.9250 | 0.8938 | 0.9058 |
| **s.e.m.** | Stytra | 0.0135 | 0.0467 | 0.0522 | 0.0482 |
| | Offline | 0.0070 | 0.0154 | 0.0428 | 0.0285 |
| | Online | 0.0088 | 0.0138 | 0.0564 | 0.0375 |
| **mean_high** | Stytra | 0.9870 | 0.9437 | 0.9783 | 0.9606 |
| | Offline | 0.9847 | 0.9518 | 0.9522 | 0.9514 |
| | Online | 0.9833 | 0.9377 | 0.9620 | 0.9495 |
| **sem_high** | Stytra | 0.0030 | 0.0112 | 0.0101 | 0.0101 |
| | Offline | 0.0028 | 0.0178 | 0.0181 | 0.0116 |
| | Online | 0.0031 | 0.0108 | 0.0162 | 0.0105 |
| **mean_low** | Stytra | 0.9375 | 0.7409 | 0.8138 | 0.7739 |
| | Offline | 0.9652 | 0.9244 | 0.8089 | 0.8608 |
| | Online | 0.9560 | 0.8995 | 0.7574 | 0.8183 |
| **sem_low** | Stytra | 0.0321 | 0.0670 | 0.1490 | 0.1046 |
| | Offline | 0.0208 | 0.0342 | 0.1085 | 0.0769 |
| | Online | 0.0244 | 0.0345 | 0.1350 | 0.0944 |

**Table S7. Within-session correlation between bout amplitude and time.** Pearson correlation (r and p) between peak-to-peak bout amplitude and time within session for each of the six testing fish (n bouts per fish) and pooled across sessions (n = 2948 bouts; pooled r = +0.2535), showing no consistent within-session amplitude trend across fish.

| Fish ID (Type) | Number of bouts | Pearson r | p-value |
|---|---|---|---|
| **241023 0101 (H)** | 496 | 0.1459 | 0.0011 |
| **241023 0202 (L)** | 595 | 0.1857 | < 0.0001 |
| **241119 0501 (H)** | 572 | 0.0371 | 0.3760 |
| **241121 0201 (L)** | 377 | 0.4540 | < 0.0001 |
| **250402 0101 (H)** | 306 | -0.4386 | < 0.0001 |
| **250402 0201 (H)** | 602 | -0.1333 | 0.0010 |
| **Pooled** | 2948 | 0.2535 | < 0.0001 |

**Table S8. Per-fish detection metrics on the free-swimming sessions.** Accuracy, precision, recall, and F1 for the threshold detector, offline model, and online model on three labeled free-swimming sessions (N = 3, 6-7 dpf, WT AB), with group means ± SEM.

| Fish ID | Method | Accuracy | Precision | Recall | F1 Score |
|---|---|---|---|---|---|
| **250318 0101** | Stytra | 0.8993 | 0.6961 | 0.9001 | 0.7851 |
| | Offline | 0.9576 | 0.8679 | 0.9349 | 0.9002 |
| | Online | 0.9429 | 0.8245 | 0.9152 | 0.8675 |
| **250318 0201** | Stytra | 0.9974 | 0.9161 | 0.9659 | 0.9403 |
| | Offline | 0.9981 | 0.9643 | 0.9476 | 0.9559 |
| | Online | 0.9976 | 0.9475 | 0.9372 | 0.9423 |
| **250318 0301** | Stytra | 0.7951 | 0.3841 | 0.9219 | 0.5423 |
| | Offline | 0.9443 | 0.7197 | 0.9450 | 0.8171 |
| | Online | 0.9321 | 0.6821 | 0.9071 | 0.7787 |
| **mean** | Stytra | 0.8973 | 0.6654 | 0.9293 | 0.7559 |
| | Offline | 0.9667 | 0.8506 | 0.9425 | 0.8911 |
| | Online | 0.9575 | 0.8180 | 0.9198 | 0.8628 |
| **s.e.m.** | Stytra | 0.0584 | 0.1543 | 0.0194 | 0.1158 |
| | Offline | 0.0162 | 0.0711 | 0.0039 | 0.0403 |
| | Online | 0.0203 | 0.0767 | 0.0090 | 0.0473 |

**Table S9. *adgrl3.1* bout duration: offline model versus best threshold method.** Two-way ANOVA main effects and Tukey post-hoc results for *adgrl3.1* KO versus WT duration under each condition, for both detectors. The offline model reaches significance on two contrasts not reached by the threshold method (CL-ON Trial 4 WT < KO; OL-ON strain main effect), with none unique to the threshold method.

| Condition | Trial Effect | | Strain Effect | | |
|---|---|---|---|---|---|
| | Offline | Threshold | Trial | Offline | Threshold |
| CL-ON | (0.0355) | (0.0481) | Main | (0.0001) | (0.0138) |
| | | | T4 | WT < KO (0.0211) | n.s. (0.5051) |
| CL-OFF | n.s. (0.5276) | n.s. (0.5980) | Main | n.s. (0.8603) | n.s. (0.3775) |
| OL-ON | n.s. (0.7855) | n.s. (0.9793) | Main | (0.0016) | n.s. (0.2171) |
| OL-OFF | n.s. (0.4904) | n.s. (0.7305) | Main | n.s. (0.3989) | n.s. (0.8081) |

**Table S10. WT/nacre/casper bout frequency: offline model versus best threshold method.** Two-way ANOVA and Tukey results for each condition, both detectors. Six contrasts differ: five reached only by the offline model and one (OL-OFF WT T1 > T4; threshold p = 0.0434, offline p = 0.1202) only by the threshold method.

| Condition | Trail Effect | | | Strain Effect | | |
|---|---|---|---|---|---|---|
| | Strain | Offline | Threshold | Trial | Offline | Threshold |
| CL-ON | Main | (<0.0001) | (<0.0001) | Main | n.s. (0.3242) | n.s. (0.4049) |
| | NCR | T2 > T4 (0.0123) | n.s. (0.1240) | T4 | WT < CSP (0.0284) | n.s. (0.0980) |
| CL-OFF | Main | (< 0.0001) | (< 0.0001) | Main | n.s. (0.2376) | n.s. (0.2297) |
| | WT | T1 > T3 (0.0216) | n.s. (0.0599) | | | |
| | NCR | T2 > T4 (0.0191) | n.s. (0.4822) | | | |
| OL-ON | Main | (<0.0001) | (<0.0001) | Main | n.s. (0.3025) | n.s. (0.2782) |
| | NCR | T1 > T4 (0.0152) | n.s. (0.1700) | | | |
| OL-OFF | Main | (<0.0001) | (<0.0001) | Main | n.s. (0.0516) | n.s. (0.0867) |
| | WT | n.s. (0.1202) | T1 > T4 (0.0434) | | | |

**Table S11. WT/nacre/casper bout duration: offline model versus best threshold method.** Two-way ANOVA and Tukey results for each condition, both detectors. Eight contrasts differ, four reached only by the offline model and four only by the threshold method.

| Condition | Trail Effect | | | Strain Effect | | |
|---|---|---|---|---|---|---|
| | **Strain** | **Offline** | **Threshold** | **Trail** | **Offline** | **Threshold** |
| CL-ON | Main | (<0.0001) | (0.0004) | Main | (0.0297) | (0.0018) |
| | | | | T4 | n.s. (0.0857) | NCR<CSP (0.0316) |
| CL-OFF | Main | n.s. (0.1023) | n.s. (0.4077) | Main | (0.0003) | (0.0001) |
| | | | | T1 | n.s. (0.0612) | WT>NCR (0.0074) |
| | | | | T2 | n.s. (0.0838) | WT>NCR (0.0110) |
| OL-ON | Main | (0.0070) | n.s. (0.2605) | Main | n.s. (0.1770) | (0.0034) |
| | NCR | T1<T3 (0.0471) | n.s. (0.1605) | T1 | n.s. (0.5601) | WT>NCR (0.0410) |
| | CSP | T1<T4 (0.0342) | n.s. (0.9477) | T4 | NCR<CSP (0.0266) | n.s. (0.2256) |
| OL-OFF | Main | n.s. (0.4214) | n.s. (0.3518) | Main | (0.0010) | (0.0002) |
| | | | | T4 | WT>NCR (0.0425) | n.s. (0.0651) |

**Table S12. WT/nacre/casper interbout interval: offline model versus best threshold method.** Two-way ANOVA and Tukey results for each condition, both detectors. Six contrasts differ: two reached only by the offline model (including the CL-ON strain × trial interaction, offline p = 0.0115, threshold p = 0.1626) and four only by the threshold method (predominantly within-casper CL-OFF progressions).

| **Condition** | **Trail Effect** | | | **Strain Effect** | | |
|---|---|---|---|---|---|---|
| | **Strain** | **Offline** | **Threshold** | **Trail** | **Offline** | **Threshold** |
| CL-ON | Main | (0.0002) | (0.0037) | Main | (0.0055) | (0.0041) |
| | Interaction | Offline: (0.0115)<br>Threshold: n.s. (0.1626) | | | | |
| CL-OFF | Main | (0.0011) | (0.0314) | Main | n.s.<br>(0.7842) | n.s.<br>(0.3643)) |
| | CSP | n.s.<br>(0.0507) | T1<T3<br>(0.0092) | T3 | n.s.<br>(0.7336) | NCR<CSP<br>(0.0461) |
| | CSP | n.s.<br>(0.0729) | T2<T3<br>(0.0217) | | | |
| OL-ON | Main | n.s.<br>(0.0955) | n.s.<br>(0.3821) | Main | n.s.<br>(0.5374) | n.s.<br>(0.3116) |
| | Interaction | Offline: n.s. (0.1232)<br>Threshold: (0.0350) | | | | |
| OL-OFF | Main | (0.0243) | n.s.<br>(0.2750) | Main | n.s.<br>(0.2473) | (0.0044) |
| | NCR | T1<T4<br>(0.0450) | n.s.<br>(>0.9999) | | | |

**Table S13. Minimum detectable effect sizes (MDES).** MDES computed at $\alpha = 0.05$ and $1 - \beta = 0.80$ for each two-way ANOVA comparison family (**Figures 4A-H**, **5**, and **S13-15**), reported as Cohen's f and partial $\eta^2$ for the strain/cohort and trial main effects and their interaction ($f = 0.2974$ to $0.5223$; partial $\eta^2 = 0.0812$ to $0.2144$). Computed in Python (statsmodels, FTestAnovaPower).

| **Comparison** | **Sample sizes** | **Test** | **Minimum detectable effect size (MDES)** |
|---|---|---|---|
| Free-swim vs Head-fixed WT<br>(Figure 4K-O) | N = 11 free,<br>N = 55 fixed | 2-way ANOVA<br>+ Fisher's LSD post-hoc | Post-hoc pairwise comparison:<br>Free vs Fixed (any stim): d = 0.9395;<br>Free vs Free across stim: d = 1.2560;<br>Fixed vs Fixed across stim: d = 0.5391 |
| *adgrl3.1*$^{-/-}$ vs WT,<br>Strain × Trial<br>(Figure 4A-H, S13) | N = 11 adgrl,<br>N = 55 WT,<br>4 trials | 2-way ANOVA<br>+ Tukey post-hoc | strain f = 0.3501, $\eta^2_p = 0.1092$;<br>trial f = 0.4195, $\eta^2_p = 0.1496$;<br>interaction f = 0.4947, $\eta^2_p = 0.1966$ |
| CL vs OL and<br>ON vs OFF in head-fixed WT<br>(Figure 4P-T) | N = 55 | 2-way ANOVA<br>+ Fisher's LSD post-hoc | Post-hoc pairwise comparison:<br>d = 0.5391 |
| Strain × Trial in<br>head-fixed WT/NCR/CSP<br>(Figure 5, S15) | N = 55 WT,<br>N = 31 NCR,<br>N = 26 CSP,<br>4 trials | 2-way ANOVA<br>+ Tukey post-hoc | strain f = 0.2974, $\eta^2_p = 0.0812$;<br>trial f = 0.3177, $\eta^2_p = 0.0917$;<br>interaction f = 0.4062, $\eta^2_p = 0.1417$ |
| Stytra-driven vs<br>online-model-driven<br>head-fixed WT,<br>Cohort × Trial<br>(Figure S14) | N = 55 Stytra,<br>N = 5 online,<br>4 trials | 2-way ANOVA<br>+ Tukey post-hoc | cohort f = 0.3678, $\eta^2_p = 0.1192$;<br>trial f = 0.4415, $\eta^2_p = 0.1631$;<br>interaction f = 0.5223, $\eta^2_p = 0.2144$ |
